# The MYCN 5′ UTR as a therapeutic target in neuroblastoma

**DOI:** 10.1101/2024.02.20.581230

**Authors:** Marina P Volegova, Lauren E Brown, Ushashi Banerjee, Ruben Dries, Bandana Sharma, Alyssa Kennedy, John A. Porco, Rani E George

**Author notes:** Center for Genomic Medicine, Mass General Brigham, Boston, MA, United States.

## Abstract

Tumor cell amplification of the MYCN transcription factor is seen in half of patients with high-risk neuroblastoma, where it functions as an oncogenic driver associated with metastatic disease and poor survival. Yet, direct targeting of MYCN has been met with little success, prompting efforts to inhibit its expression at multiple levels. MYCN-amplified neuroblastoma cells have an increased requirement for protein synthesis to meet the overwhelming transcriptional burden imposed by oncogenic MYCN. Here, we take advantage of this vulnerability to interrogate the therapeutic potential of inhibiting the activity of the eukaryotic translation initiation factor 4A1 (eIF4A1), an RNA-helicase responsible for resolving structural barriers such as polypurine preponderance within 5′ untranslated regions (UTRs). We observed that eIF4A1 is a key regulator of transcript-specific mRNA recruitment in MYCN-overexpressing neuroblastomas and MYCN-associated transcripts rank highly in polypurine-rich 5′ UTR sequences, the majority of which have critical roles in cell proliferation. Using CMLD012824, a novel synthetic amidino-rocaglate (ADR) derivative, we demonstrate selectively increased eIF4A1 affinity for polypurine-rich 5′ UTRs, including the MYCN mRNA, leading to translation inhibition and cytotoxicity in human neuroblastoma cell lines and animal models. Through ribosome profiling and PAR-CLIP analysis, we show that ADR-mediated clamping of eIF4A1 onto mRNA spans the full lengths of target transcripts, whereas translational inhibition is mediated selectively through 5′ UTR binding. Both cap-dependent and cap-independent translation of MYCN are disrupted, pointing to the ability of CMLD012824 to disrupt non-canonical translation initiation. Our studies provide insights into the functional role of eIF4A1 in meeting the increased protein synthesis demands of MYCN-amplified neuroblastoma and suggest that its disruption may be therapeutically beneficial in this disease.

## Introduction

Neuroblastoma (NB), a tumor of the sympathetic nervous system, accounts for 8-10% of all childhood cancers and has a survival rate of < 50% in patients with high-risk disease ^1^. Nearly half of all high-risk tumors harbor amplification of MYCN, an oncogenic driver encoding a member of the MYC family of transcription factors which is significantly associated with aggressive disease and fatal relapse ^2^. As with other MYC family members, MYCN is considered “undruggable”, largely due to the lack of drug binding surfaces on its helix-loop-helix structure ^3,4^, prompting investigations into disrupting the expression of MYCN and its downstream targets for therapeutic purposes ^5–8^. Recent studies of translational control in MYC-driven cancers have identified components of the mRNA translation machinery as major drivers of oncogenesis ^9,10^. While protein synthesis is a feature of all cells, cancers driven by oncogenic transcription factors, such as MYCN-amplified NB have a corresponding need for translational upregulation to meet the overwhelming transcriptional burden imposed by oncogenic MYCN. Indeed, MYCN-amplified NB cells exhibit significant upregulation and dependence on both ribosome biogenesis and protein translation ^11,12^ suggesting that these processes could be disrupted for therapeutic benefit.

Much of translation regulation in eukaryotic cells occurs at the initiation step, a process mediated by the eukaryotic initiation factor 4F (eIF4F) complex, composed of eIF4A, the ATP-dependent DEAD-box RNA helicase that is crucial for unwinding 5′ untranslated region (5′ UTR) secondary structures and preparing a clear path for ribosome scanning, as well as the cap binding protein eIF4E and the scaffolding protein eIF4G ^13^. Upon binding to the mRNA cap, the eIF4F complex remodels the 5′ UTR and recruits the 43S ribosome pre-initiation complex (PIC) ^13,14^. The PIC then scans the 5′ UTR for an initiation codon to start the translation process. Hence, mRNAs must compete for access to the eIF4F complex, and structural barriers within their 5′ UTRs can impact their reliance on eIF4F and its ability to recruit or alter the scanning efficiency of the PIC ^15,16^. This is especially true of oncogenic mRNAs such as that of MYC whose complex 5′ UTR secondary structures render them heavily dependent on the eIF4A helicase for translation ^17^. The ribonucleotide composition of the 5′ UTR is primarily responsible for this effect, where stem loop formation, GC and AG content, and G-quadruplexes all play a role in negatively impacting the speed of translation initiation ^18–20^.

Due to its critical role in gene expression, translation initiation is frequently commandeered by oncogenic drivers to regulate the expression of growth-promoting genes, and thus has emerged as an attractive therapeutic target ^9,14,21,22^. However, while an abundance of compounds capable of disrupting translation initiation exist ^23,24–26^ only a member of the rocaglate family, zotatifin (eFT226) ^27,28^ has entered clinical trials to date. Rocaglates are naturally occurring compounds containing a common cyclopenta[*b*]benzofuran core and together with their synthetic analogs, are highly potent protein synthesis inhibitors. These compounds repress translation by causing eIF4A1 (the primary eIF4A homolog) to preferentially clamp onto polypurine-rich sequences in the 5′ UTRs of mRNAs, thereby blocking ribosome scanning ^29,30^. Such activity of rocaglates on complex 5′ UTRs provides a selective therapeutic advantage in cancer cells due to the polypurine-rich 5′ leaders of oncogenic and cell cycle-regulating mRNAs ^26^. The synthetic rocaglate hydroxamate, CR-1-31-B, has been tested in several cancers including NB^17,24,31^, although whether it exhibits selective transcript-specific effects without inducing systemic toxicity is not fully understood. Here, we employ a new class of synthetic rocaglate analogs, amidino-rocaglates (ADRs)^32,33^, that target eIF4A1 with higher specificity and selectivity to investigate translation factor dependence and inhibition of translation initiation in MYCN-amplified NB.

## Results

### eIF4A1 expression is enriched in MYCN-amplified NB

To determine the therapeutic potential of inhibiting protein translation in NB, we first examined the expression of translation initiation factors in primary human tumors through analysis of an RNA-sequencing data set comprising both MYCN-amplified and nonamplified tumors (n = 498; MYCN amplified = 92; GSE62564). Based on the annotated MYCN amplification status of the dataset, higher expression of several factors was observed in MYCN-amplified compared to MYCN-nonamplified tumors, most significantly of mRNAs that comprise the eIF4F complex (eIF4A1, eIF4E, eIF4G1) (Fig. S1A, 1B). On analyzing the entire tumor cohort based on MYCN expression levels (from lowest to highest), we again observed a positive correlation between MYCN expression and that of eIF4F members, with tumors having higher MYCN expression exhibiting higher expression of these factors (Fig. 1A). The strongest correlation was noted between the MYCN transcript itself and that of eIF4A1, and to a lesser extent, with that of eIF4E and eIF4G1 (Fig. 1B). Interestingly, eIF4F complex expression was not significantly associated with that of c-MYC overexpression, which has been reported in a subset (∼10%) of MYCN-nonamplified NBs ^34^ (Fig. 1C, 1D). Pairwise correlation analysis of the MYCN-amplified tumor subset (n = 92) confirmed the positive correlation between higher MYCN and eIF4A1 and eIF4G1 transcript levels, but not that of eIF4E (Fig. 1E). Contrastingly, there was a lack of correlation between c-MYC and eIF4A1 expression in the MYCN-nonamplified tumor subset (n = 401) (Fig. S1C). In addition, analysis of our previously published MYCN chromatin immunoprecipitation and high-throughput sequencing (CHIP-seq) data in MYCN-amplified NB cells ^35^ (GSE103084) revealed that MYCN binds to the promoters of genes encoding the eIF4F complex, but at much higher levels for *EIF4A1* compared to *EIF4E* and *EIF4G1* (Fig. 1F). Together, these results suggest that eIF4A1 may play a prominent role in MYCN-driven translation and that its inhibition may be deleterious to MYCN-amplified NB.

**Figure 1.**
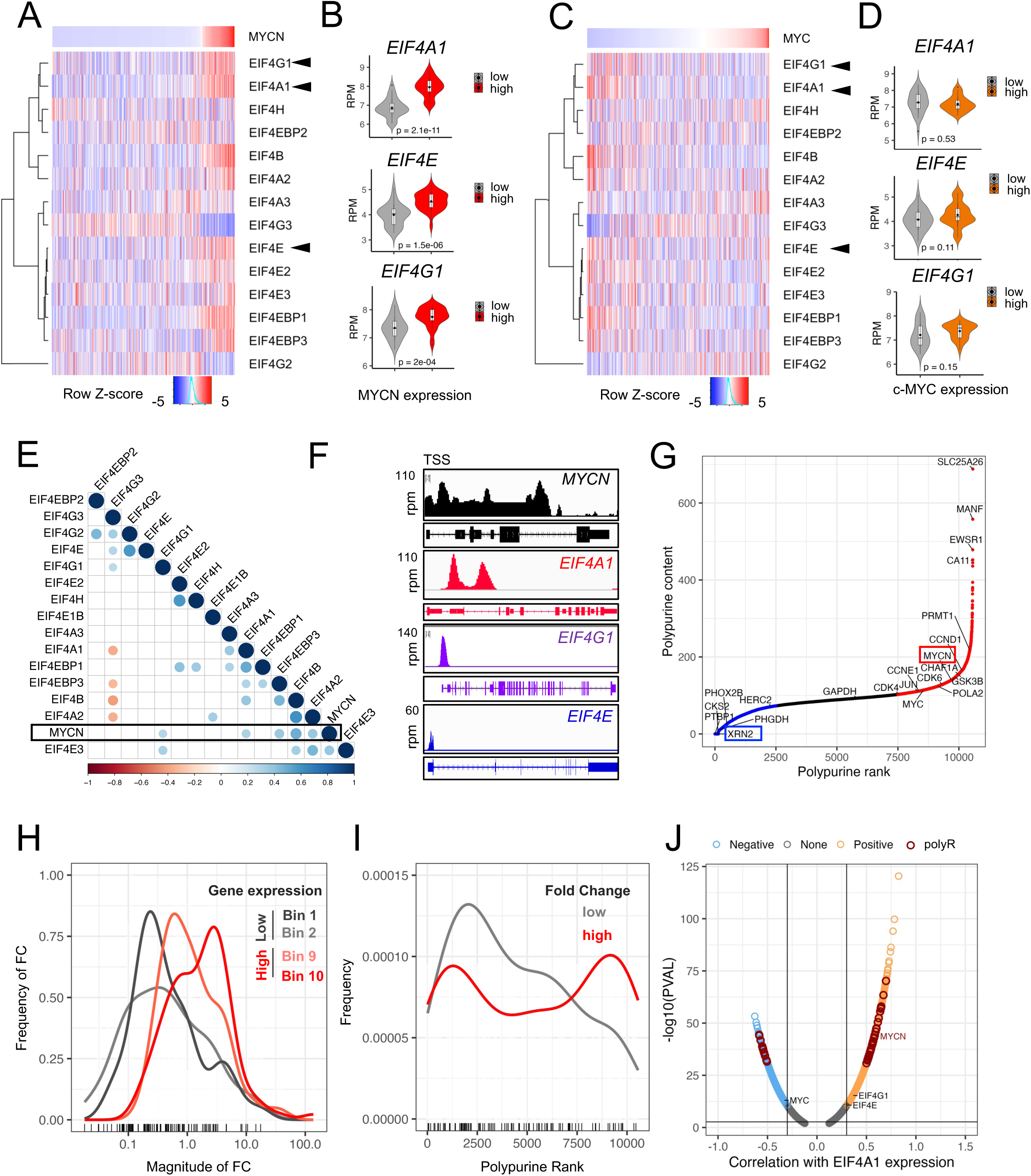
MYCN-amplified NBs exhibit upregulation of translation initiation factors and are enriched for mRNAs with polypurine-rich 5’ UTRs. **A.** Hierarchical clustering of translation initiation factor gene expression in primary NB tumors (n=498, GSE62564), ranked by MYCN expression. z-score = mean ± S.D. Bar above heatmap represents corresponding MYCN expression level in log reads per million (log_2_ RPM). **B**. Violin plots showing expression of the indicated initiation factors in tumors with lowest and highest MYCN expression levels, as depicted in **A** (n=30 each). Box plots within the violin plots defined by center lines (medians), box limits (the interquartile range between 25^th^ and 75^th^ percentiles), whiskers (minima and maxima;). Significance determined by Student’s t-test. **C.** Hierarchical clustering of the tumors in **A**, ranked by c-MYC expression. z-score = mean ± S.D. Bar above heatmap represents corresponding c-MYC expression level (RPM). **D.** Violin plots depicting the expression of the indicated initiation factors in primary tumors in **C** with the highest and lowest c-MYC (n = 30) expression levels. Box plots within the violin plots defined as in **B**. **E.** Correlogram of MYCN and translation initiation factor gene expression in MYCN-amplified primary tumors (n = 92, GSE62564). Circles represent Spearman’s rank correlation coefficients, *P* <0.01. Color code represents positive correlations in blue, negative correlations in red. **F.** ChIP-seq profiles of MYCN binding at the *indicated* gene loci in Kelly NB cells. The x-axis shows genomic position and the y-axis the signal of MYCN binding in units of reads per million per base pair (rpm). **G**. Ranking of mRNAs expressed in primary NBs based on polypurine content [calculated as the number of polypurine ([AG]_n_) motifs in the 5′ UTRs of their corresponding mRNAs] and plotted in order of increasing polypurine rank from lowest (1) to highest (10570). Upregulated mRNAs (>2-fold change) are shown in red, downregulated mRNAs in blue, no change in black, with fold change calculated as changes in expression of highly variable genes (Student’s t-test, *P* < 0.01). **H.** Fold change distributions of highly variable genes (HVGs) in tumors with lowest [bottom 10% (bins 1 and 2)] and highest [top 10% (bins 9 and 10)] MYCN expression levels (n=30 each, GSE62564) (Student’s t-test, *P* < 0.01). X-axis shows the magnitude and y-axis shows the frequency of fold change (FC) in expression. **I.** Polypurine rank distribution of the highly variable upregulated genes [high (>5-fold change), red; low (2-fold change), gray] (Student’s t-test, *P* < 0.01). **J.** Volcano plot of genes correlated with eIF4A1 in primary tumors (n=498, False discovery rate (FDR) < 0.05). Gold, positive; blue, negative correlation; red, genes that ranked in the top 25% of polypurine content.

As the helicase responsible for facilitating ribosome progression along mRNA, eIF4A1 must unwind complex structures that serve as roadblocks to the PIC scanning mechanism, including 5′ UTR polypurine stretches ([AG]_n_, which are targets of rocaglate inhibitors ^36^. To assess the targetability of MYCN-amplified NB with rocaglates, we analyzed the 5′ UTR polypurine content of the transcripts expressed in the 498 primary tumor data set (GSE62564) by quantifying sequential polypurines in the corresponding 5′ UTR regions of all expressed transcripts and ranking these from lowest (polypurine-poor, rank = 1) to highest (polypurine-rich, rank = 10570) after normalizing to 5′ UTR length (Fig. 1G). The MYCN 5′ UTR itself ranked highly in polypurine content (rank = 9689, 93^rd^ percentile); by contrast, the c-MYC 5′ UTR had a relatively lower polypurine content (rank = 8552, 80^th^ percentile), in keeping with the lack of sequence homology between the two 5′ UTRs^37^. Transcripts encoding genes with major roles in cell proliferation - CCND1, CCNE1 and CDK4/6 - were also represented among the top polypurine-rich group (Fig. 1G). Next, we sought to understand the extent of 5′ UTR polypurine content of transcripts that were differentially expressed between MYCN-amplified and -nonamplified tumors on the premise that these mRNAs would be the most biologically relevant. We first identified the highly variably expressed genes in all the tumors by arraying and binning all transcripts by expression level and calculating the variance coefficient as previously described ^38^ (Fig. S1D). We next identified the top and bottom 30 tumors ranked by MYCN expression levels (MYCN-amplified and -nonamplified subsets respectively) and calculated the fold-change in expression of the highly variable transcripts within these tumors (n = 524). Among these transcripts, 30% were upregulated (n = 162/524) and 60% were downregulated (n=317/524) in the top MYCN-expressing (MYCN-amplified) compared with the bottom (MYCN-nonamplified) tumors), with, unsurprisingly, MYCN emerging as the most significantly upregulated transcript (Fig. S1E). Among the highly variable mRNAs, those with the highest expression were more likely to be upregulated in MYCN-amplified vs. -nonamplified tumors (Fig. 1H, S1F). To highlight the differences in polypurine content observed at the extreme ends of the dataset, we next analyzed these most differentially expressed genes based on polypurine ranking and observed that the most upregulated genes were enriched for high polypurine content (Figs. 1I, S1H). The highly expressed polypurine-rich mRNAs specific to MYCN-amplified NBs were functionally enriched in key cellular processes, such as the G2/M checkpoint and RNA processing and quality control, as well as MYC targets (Fig. S1I) suggesting that their inhibition would negatively impact cell proliferation. Finally, within the primary tumor data set, we observed that transcripts with polypurine-rich 5′ UTRs, including that of MYCN, were significantly overrepresented within transcripts that were positively correlated with eIF4A1 overexpression (surpassing even eIF4E and eIF4G1) compared to those that were negatively correlated (Fig.1J). Together, these analyses demonstrate that MYCN-amplified tumors are enriched in transcripts with polypurine-rich 5′ UTRs that are highly correlated with eIF4A1 expression and suggest that they could be amenable to rocaglate-mediated inhibition.

### eIF4A1 inhibition is selectively cytotoxic to MYCN-amplified NB

To identify a rocaglate derivative with high specificity and selectivity for eIF4A1, we screened a library (n = 42) of synthetic rocaglate analogs against a panel of established and patient-derived xenograft (PDX) human NB cell lines and identified compound **CMLD012824** (hereafter referred to as “**ADR-824**” in Figures), as highly potent (Figs. 2A, S2A). CMLD012824 is a member of the amidino rocaglate (ADR) series of compounds, which differ structurally from other rocaglates by the addition of a 2-imidazoline or cyclic amidine ring. The chiral, racemic version of this compound (CMLD012612), which includes the non-bioactive enantiomer, was previously found to inhibit lymphoma growth in mice in combination with doxorubicin, but as a single agent had no effect on tumor-free survival ^39^. CMLD012824 is the pure form of the bioactive enantiomer, which had previously been found to be cytotoxic in one breast cancer cell line ^32^. CMLD012824 exhibited relatively higher potency against MYCN-amplified NB cells, with a half maximal inhibitory concentration (IC_50_) in the sub-nM range compared to MYCN-nonamplified or non-transformed cells (Fig. 2A). MYCN-amplified cells underwent dose-independent apoptosis and loss of membrane integrity within an hour of treatment, while nonamplified cells reached peak apoptotic response only at 24 hours (Figs. 2B, Fig. S2B). Additionally, CMLD012824 led to both G1 and G2 cell cycle arrest in MYCN-amplified cells, but primarily G2 arrest in MYCN-nonamplified cells (Figs. 2C, S2C). Importantly, HEK293 non-transformed cells showed no cycling defects at similar treatment conditions suggesting that the cytotoxic effects of the CMLD012824 ADR derivative may be selective for cancer cells (Figs. 2C, S2C). Consistent with the differential effects on the cell cycle, the decreased expression of regulatory proteins (CCND1, CCNE1, CDK4), was observed at lower doses in MYCN-amplified versus nonamplified cells (Fig. 2D). In keeping with its putative mode of action, CMLD012824 did not affect total eIF4A1 protein levels (Fig. 2D). Finally, to assess the global impact of CMLD012824 on protein synthesis, we performed metabolic labeling of nascent proteins in MYCN-amplified, nonamplified and non-transformed cells. In comparison with the promiscuous protein synthesis inhibitor cycloheximide, which abrogated protein synthesis in all three cell types, CMLD012824 preferentially inhibited protein synthesis in MYCN-amplified NB cells and less so in MYCN-nonamplified and non-transformed cells (Fig. 2E). Together, these results illustrate the divergent cellular responses elicited by CMLD012824 and suggest that this ADR analog may be selectively toxic to malignant cells and, in particular, to MYCN-amplified NB cells.

**Figure 2.**
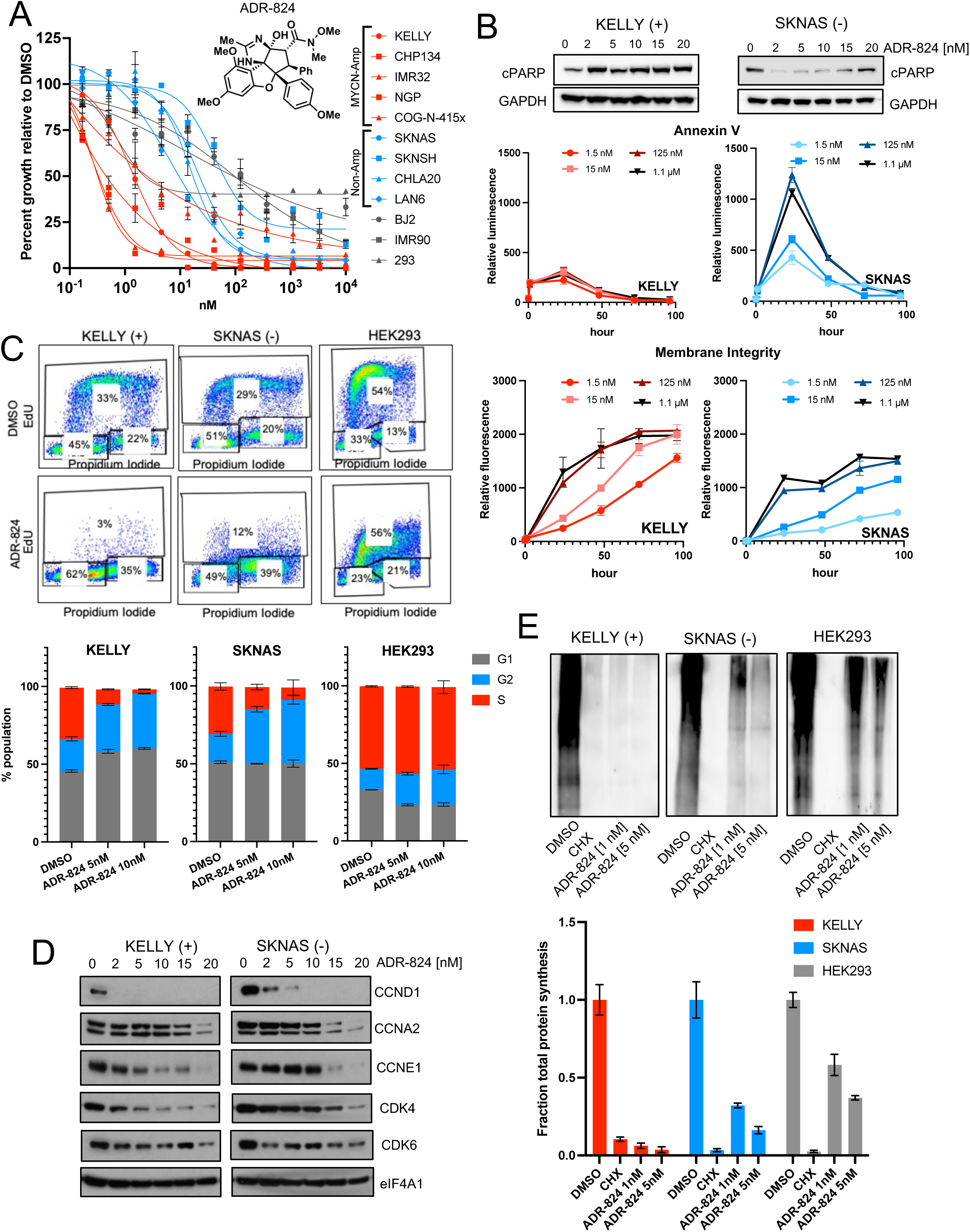
The CMLD012824 ADR exhibits differential cytotoxicity in NB cells. **A.** Cell viability of MYCN amplified (red), non-amplified (blue) human NB and non-transformed (gray) cells, treated with varying concentrations of CMLD012824 (ADR-824) for 72 h. Percent cell viability relative to DMSO is shown. Data represent mean ± S.D., n = 3 replicates. Inset: chemical structure of ADR-824. **B.** Upper, western blot (WB) analysis of PARP cleavage, GAPDH loading control; Middle, annexin V; Lower, membrane integrity analyses in MYCN-amplified (Kelly) and nonamplified (SK-N-AS) cells exposed to ADR-824 at the indicated doses. Data represent mean ± S.D., n = 3 replicates. **C**. Flow cytometry analysis of propidium iodide and EdU incorporation in the indicated NB (Kelly, SK-N-AS) and non-transformed (293) cells 24 h post exposure to ADR-824 (10 nM). Bottom, quantification of mean ± S.D. n = 3 biological replicates. **D.** WB analysis of cell cycle markers in the indicated cells 24 h after varying doses of ADR-824. eIF4A1 represents the loading control. **E.** Metabolic labeling with L-azidohomoalanine (AHA) of nascent protein synthesis in the indicated cells exposed to ADR-824 (1 nM, 5 nM), cycloheximide or DMSO for 1 hour. Blot represents n=3 biological replicates. Bottom, quantification of mean ± S.D. of 2 replicates.

### ADR-mediated eIF4A1 inhibition selectively decreases MYCN translation

Given the relatively high polypurine content of its 5′ UTR (Fig. 1G), we predicted that the MYCN transcript would be especially sensitive to CMLD012824-mediated translation inhibition. Indeed, treatment of MYCN-amplified NB cells led to a complete loss of MYCN protein signal on immunofluorescence microscopy compared to that of PTBP1, a 5′ UTR polypurine-poor control (Figs. 3A, 1G). Concomitantly, a dose-dependent decrease in MYCN protein levels was seen in MYCN-amplified NB cells (Fig. 3B). Meanwhile c-MYC protein levels in MYCN-nonamplified NB cells were less affected, consistent with the lower polypurine content ranking of the c-MYC 5′ UTR (Fig. 3B). Neither the MYCN-amplified nor nonamplified cells were able to increase eIF4A1 protein levels to compensate for the inhibitory effect (Fig. 3C). To confirm whether the sensitivity of specific proteins to ADR inhibition could be predicted based on the polypurine content of their respective mRNAs, we assessed the effect of CMLD012824 on the translation of the polypurine-rich MYCN and the polypurine-poor XRN2 proteins (Fig. 1G) in comparison to the global protein synthesis inhibition induced by cycloheximide. While cycloheximide led to reduced levels of both proteins, CMLD012824 caused loss only of MYCN and not XRN2 levels (Fig. 3D). MYCN protein loss was sustained despite compensatory transcriptional upregulation of the mRNA (Fig. 3E). Next, we determined whether transcription or protein degradation contributed to the effects of CMLD01284. As expected, the proteasomal inhibitor, MG132, alone or in combination with the transcription inhibitor actinomycin D led to a slight increase in MYCN protein levels, likely due to inhibition of degradation and translation of accumulated RNA, respectively (Fig. 3F). Both CMLD01284 and MG132 individually or together did not significantly affect MYCN levels, indicating that the function of CMLD01284 is not proteasome-dependent. On the other hand, while actinomycin D alone did not substantially affect MYCN levels, the addition of CMLD01284 led to a striking reduction, which was rescued by MG132. These results together point to inhibition of translation as the primary function of CMLD01284. Finally, we determined whether the loss of MYCN with CMLD012824 treatment also affected its function as a DNA-binding transcription factor as shown in Fig. 1F. Chromatin immunoprecipitation of the MYCN protein followed by RT-qPCR (ChIP-qPCR) showed a decrease in MYCN occupancy at the promoters of various genes, including MYCN itself and known targets EIF4A1 (Fig. 1F), TP53, and AURKA ^40,41^ (Fig. 3G). In contrast, the polypurine-poor PHOX2B transcription factor (Fig. 1G) showed no change in occupancy at its target promoters following CMLD012824 treatment (Fig. 3G). These findings allow us to conclude that impaired translation of MYCN is one of the main mechanisms through which CMLD012824 exerts it cytotoxic effects in MYCN-amplified NB cells.

**Figure 3.**
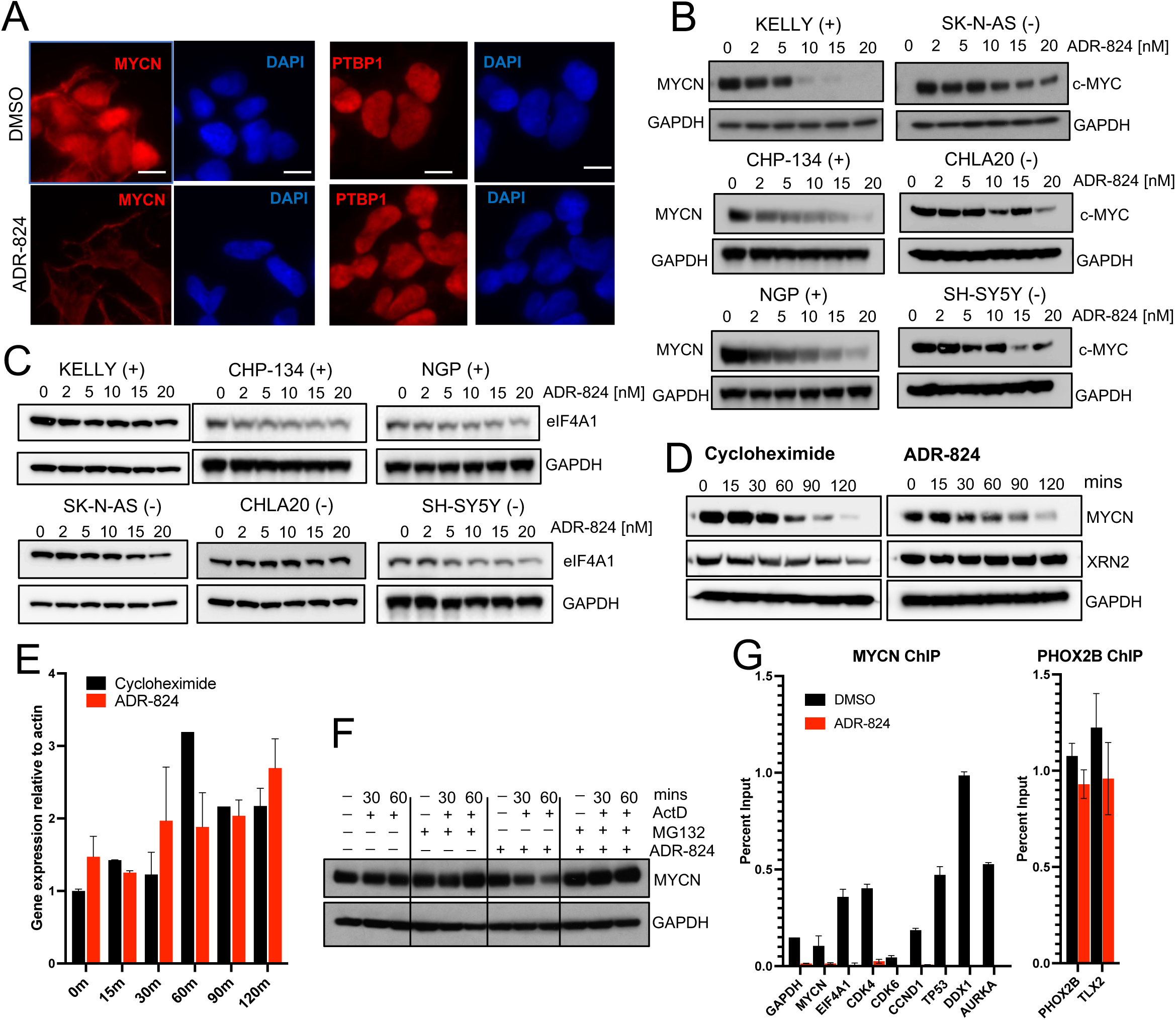
ADR-mediated inhibition of eIF4A1 leads to MYCN protein degradation. **A.** Immunofluorescence images of the MYCN protein in MYCN-amplified (Kelly) cells at 1 h post ADR-824 (10 nM) treatment. PTBP1 serves as a polypurine-poor control. DAPI nuclear stain in blue. scale bar,10 µm. **B.** WB analysis of MYCN and c-MYC expression at 4 h post ADR-824 treatment at the indicated doses in MYCN-amplified (+) and nonamplified (-) NB cell lines. **C.** WB analysis of eIF4A1 protein levels at 4 h post ADR-824 treatment at the indicated doses in MYCN-amplified (+) and non-amplified (-) NB cell lines. **D**. WB analysis of MYCN expression in MYCN-amplified (Kelly) cells treated with CHX (10 µg/mL) or ADR-824 (10 nM) for the indicated times. XRN2 is used as a polypurine-poor control. **E.** RT-qPCR analysis of MYCN mRNA levels in cells treated with cycloheximide or ADR-824 as in **D**. Data represent mean ± S.D., n = 2 replicates. **F.** WB analysis of MYCN expression in MYCN-amplified NB cells treated with actinomycin D (actD, 1 µg/mL; 30 or 60 minutes (min) with or without MG132 (100 µM) or ADR-824 (10 nM). GAPDH serves as loading control for **B, C, D** and **E. G.** ChIP-PCR analysis of MYCN and PHOX2B at the promoters of the indicated genes in MYCN-amplified NB cells under DMSO- and ADR-824-(10 nM) treated conditions. Percent binding relative to input signal and IgG control is shown. Data represent mean ± S.D., n = 3 replicates.

### CMLD012824 disrupts the translation of long and polypurine-rich 5′ UTRs

We next sought to understand whether the cytotoxicity of CMLD012824 in NB cells was due to the purported effect of rocaglates to inhibit active translation by decreasing mRNA translation efficiency ^36^. We therefore analyzed the changes in ribosome occupancy on mRNA transcripts through ribosome profiling (Ribo-seq) ^42^ of MYCN-amplified (Kelly) and nonamplified (SK-N-AS) NB cells following a 1-hr. exposure to CMLD012824. Sequencing reads of ribosome-protected fragments normalized to total RNA sequences were used to define translational efficiency as previously described^43^. Rather than a global downregulation of protein synthesis, CMLD012824 led to differential translation in both cell types compared to DMSO-treated cells. While significant decreases in translational efficiencies (33%; 1841/5621) were observed in MYCN-amplified cells, an *increase* in translation efficiencies was also noted (26%; 1451/5621) (>1.5-fold change in either direction) (Fig. 4A). Similar, but more modest numbers of differentially translated mRNAs were seen in MYCN-nonamplified cells (downregulated, 25%, 1535/6053; upregulated, 16%, 950/6053) (Fig. S3A). Downregulated mRNAs that overlapped between the MYCN-amplified and nonamplified cells (n = 994) were enriched for major proliferative and signaling processes such as WNT, NGF TGF-beta, PDGF and Notch pathways (Figs. S3B, S3C). The uniquely downregulated in MYCN-amplified cells (46%, 847/1841) were enriched for RNA polymerase II sequence-specific DNA binding and transcription regulation. Those similarly affected in MYCN-nonamplified cells (36%, 541/1535) also involved the same processes (with non-overlapping mRNAs), although the extent of differential expression varied, with effects being more significant in MYCN amplified cells (Fig. S3D). The uniquely upregulated mRNAs in MYCN-amplified cells were enriched for RNA-binding factors, such as the RNA helicase DDX52, nuclear RNA-binding protein TDP43, and initiation factor eIF1 (Fig. S3E, 4B), likely as a compensatory response to translation inhibition.

**Figure 4.**
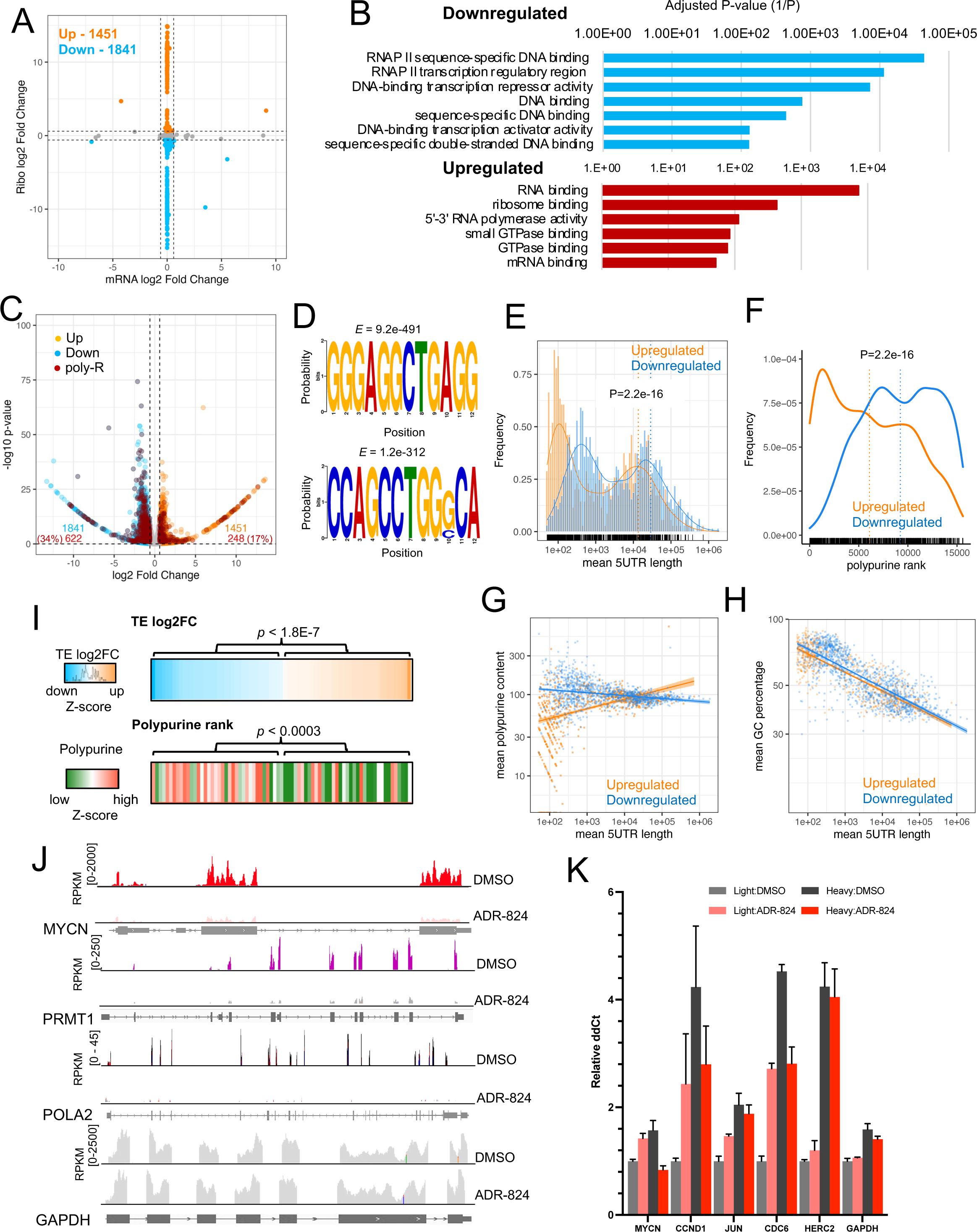
ADR-824 leads to selective translation repression of long, polypurine-rich mRNAs in NB cells. All the panels refer to ADR-824-treated MYCN-amplified (Kelly) NB cells (10 nM x 1 h). **A.** Scatter plot of total vs. ribosome-associated mRNA changes in ADR-824-treated cells, showing log_2_ fold changes as determined by Anota2seq (*P* <0.1) and normalized to synthetic RNA control (ERCC92) (see Methods). Dotted black lines indicate cut-offs of 1.5-fold change in total mRNA (x-axis) and ribosome occupancies (y-axis) of ADR-824-treated biological replicates compared to vehicle controls (n=3 each). Data points are color coded according to ribosome occupancy, with corresponding total numbers shown on the plot (upregulated, gold; downregulated, blue). **B.** Functional enrichment of unique downregulated (top, blue) and upregulated (bottom, red) transcript sets in MYCN-amplified (Kelly) cells [Down: >1.5-fold decrease, Up: >1.5-fold increase (*P* <0.1)]. Axis depicts log of inverse P-value derived from Fisher’s exact test in Enrichr. **C**. Volcano plot of translationally regulated mRNAs in DMSO- vs. ADR-824-treated cells, as determined by Anota2seq (>1.5-fold change, *P* <0.1). Polypurine-rich (top 25%) mRNAs are highlighted in red. **D.** Motif enrichment analysis of the top motifs in the downregulated mRNA subset, trained against a background list of unregulated transcripts. Statistical significance (E-values) derived from Meme. **E.** 5′ UTR length distribution of translationally regulated transcripts. (*P* <2.2e-16; up- vs. down-regulated mRNAs, >2-fold change, Student’s t-test). **F**. Polypurine rank distribution of translationally regulated transcripts. (*P* <2.2e-16; Student’s t-test). **G, H.** Scatter plots of polypurine (**G**) and GC content (**H**) changes by 5′ UTR length in upregulated (gold) versus downregulated (blue) mRNAs. Loess regression analysis is shown in corresponding colors, shaded regions represent 95% confidence intervals. **I.** Heatmaps of polypurine ranking and translational efficiency (TE) changes (n = 76; P <0.1, Anota2seq) of MYCN-regulated target genes following ADR-824 treatment compared to control cells. Z-score = mean ± S.D. P-values above heatmaps correspond to Student’s t-test of TE and polypurine rank between downregulated and upregulated genes. **J.** Ribosome occupancy profiles of the polypurine-rich *MYCN-regulated genes* in MYCN-amplified (Kelly) cells. GAPDH serves as an unaffected polypurine-poor control. Ribosome profiling signal in units of reads per kilobase per million (RPKM). **K.** RT-qPCR analysis of the indicated mRNA distributions in polysome fractions after DMSO or ADR-824 treatment in MYCN-amplified (Kelly) NB cells, pooled according to polysome occupancy. Light: 1-3 polysomes; heavy: 4+ polysomes. Signal was calculated by 2^-ΔΔCt method, normalized to total RNA in gradient and GAPDH controls. Data represent mean ± S.D., n = 3 replicates.

Consistent with the affinity of ADRs for polypurine sequences, 34% (622/1841) of the translationally downregulated mRNAs in MYCN-amplified cells possessed 5′ UTRs ranking in the top quartile of polypurine content, compared with 17% (248/1451) of upregulated mRNAs (Fig. 4C). Comparable percentages of polypurine-rich 5′ UTRs were observed in MYCN-nonamplified cells (33% of downregulated, 19% of upregulated) (Fig. S3F). Among the downregulated mRNAs in both cell types, the most significantly enriched motifs included short polypurine sequences (4-6 nucleotides) interspersed with CT nucleotides [GGGAGGCTGAGG], although also observed were highly significant motifs containing pyrimidine pairs and triplets (CC, CT, CCC, CCT) (Fig. 4D, S3G), suggesting that the mRNA transcript specificity of CMLD012824 is not exclusive to purely polypurine-rich motifs. Therefore, we questioned whether polypurine content alone was the defining characteristic of ADR-sensitive mRNAs, or whether 5′ UTR length also contributed to CMLD012824-mediated inhibition. Notably, downregulated mRNAs tended to have significantly longer 5′ UTRs (nt > 500) compared with upregulated mRNAs, which were instead enriched in shorter 5′ UTRs (nt < 200) (Fig. 4E, S4A). Moreover, the upregulated transcripts were not only relatively deficient in polypurine-rich 5′ UTRs (Fig. 4C), but in addition, were amongst the lowest-ranking polypurine-poor 5′ UTRs (Fig. 4F, S4B). Among the downregulated mRNAs, even those with short 5′ UTRs were polypurine-rich, whereas the upregulated mRNAs had a preponderance of 5′ UTRs that were both short and polypurine-poor (Figs. 4G, S4C). In contrast to polypurine content, there was no significant difference in GC content in the differentially translated genes in either cell type (Fig. 4H, S4D), further demonstrating that polypurine content and 5′ UTR length together are the main determinants of mRNA sensitivity to CMLD012824.

NB cell state is driven by a unique landscape of super-enhancers (SE), with the top SE being associated with MYCN itself ^5^. We analyzed the impact of CMLD012824 on the translational efficiencies of SE-associated genes in MYCN-amplified cells. Among the transcripts that were downregulated with CMLD012824 treatment, 7% were associated with SEs (n = 127/1841), 43% of which were enriched for polypurine-rich 5′ UTRs (n = 55/127), including MYCN (Fig. S4E). We also examined the effect of CMLD012824 on a 157-gene MYCN target signature previously defined from 88 primary NB tumors ^44^. Translationally downregulated MYCN target genes were significantly enriched for high polypurine content compared to those that were translationally upregulated (Fig. 4I). CMLD012824 treatment led to significant decreases in ribosome occupancies at MYCN and other polypurine-rich MYCN-target mRNAs identified in this data set, including PRMT1 and POLA2 (Fig. 4J). To support our ribo-seq findings, we used polysome gradient fractionation to directly examine the changes that occur in ribosome occupancy upon CMLD012824 treatment. In MYCN-amplified NB cells, we observed a shift from heavy (4+ ribosomes) to light (1-3 ribosomes) polysomes on polypurine-rich mRNAs (MYCN, CCND1, JUN, CDC6), confirming downregulation of their translational efficiencies by CMLD012824 compared to polypurine-poor and upregulated mRNAs that were unaffected (Figs. 4K, S4F, S4G). Thus, a significant proportion of genes that are associated with the deregulated MYCN cell state are impacted by CMLD012824, thereby severely crippling the proliferative feedback loops in MYCN-amplified NB.

### CMLD012824 results in promiscuous eIF4A1 clamping along sensitive mRNAs

The rocaglate series of compounds exert their effects on translation primarily by causing eIF4A1 to clamp onto the 5′ UTRs of polypurine-rich mRNAs, thereby preventing ribosome scanning ^36^. Thus, CMLD012824 would be expected to selectively increase the association of eIF4A1 with sensitive endogenous mRNAs. Indeed, RNA immunoprecipitation and quantitative PCR (RIP-qPCR) analysis of CMLD012824-treated MYCN-amplified NB cells revealed enrichment of eIF4A1 binding to several candidate polypurine-rich mRNAs, including MYCN, relative to polypurine-poor mRNAs (Fig. 5A, S5A). A similar enrichment pattern was observed in MYCN-nonamplified NB cells also, in keeping with the predicted mode of action of CMLD012824 (Fig. S5B).

**Figure 5.**
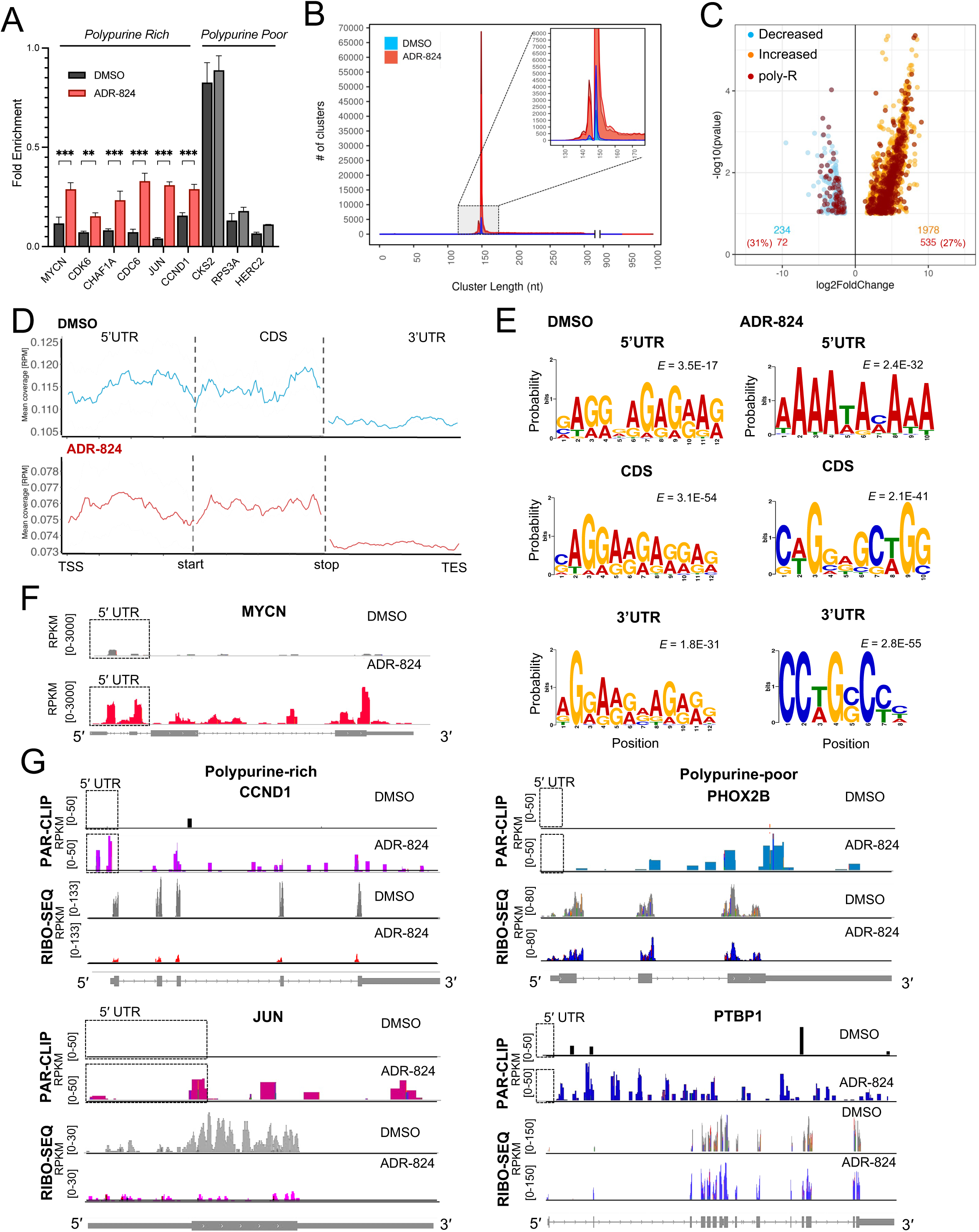
ADR-824 augments mRNA binding of eIF4A1 along the full length of mRNAs. **A.** RT-qPCR analysis of 5′ UTR polypurine-rich and -poor mRNAs bound to endogenous eIF4A1 protein immunoprecipitated from MYCN-amplified (Kelly) cell lysates following treatment with DMSO or ADR-824 (10 nM x 1 h). Data represent mean ± S.D., n = 3 replicates. **p value < 0.001, ***p value < 0.0001, Student’s t-test. **B**. Length distribution of eIF4A1 PAR-CLIP clusters. Data represent consensus clusters of two biological replicates per condition. **C**. Volcano plot of the relative changes in binding (>1.5-fold change) of eIF4A1-bound mRNAs in cells exposed to ADR- 824 or DMSO as in **A**. Statistically significant total (gold, increased; blue, decreased) and polypurine-rich (red, top 25%) mRNAs shown on graph (*P* < 0.1, Anota2seq). **D.** Metagene analysis of eIF4A1-bound clusters along the indicated mRNA regions in DMSO and ADR-824-treated cells. Data represent mean coverage (RPM), n = 2 biological replicates. **E.** Top motifs identified in eIF4A1 clusters that map to the indicated mRNA regions in DMSO- and ADR-824-treated cells. E-values adjusted to motif frequency are shown. **F**. Representative tracks of eIF4A1 binding on the MYCN mRNA from DMSO- and ADR-824-treated cells, signal in units of reads per kilobase per million (RPKM). **G.** Representative tracks of eIF4A1 binding (PAR-CLIP) and ribosome occupancy (RIBO-SEQ) profiles of polypurine-rich and -poor mRNAs from ADR-824-treated cells. Black boxes outline 5′ UTR regions.

To gain a better understanding of eIF4A1 binding across the transcriptome of MYCN-amplified NB cells, we isolated transcripts labeled with UV light-activatable 4-thiouridine that co-purified with endogenous eIF4A1 via photo-activatable ribonucleoside-enhanced cross-linking and immunoprecipitation (PAR-CLIP)^45^ in COG-N-415x MYCN-amplified patient-derived xenograft (PDX) NB tumor cells exposed to CMLD012824 (Fig. S5C). Sequencing reads of cDNA libraries generated from eIF4A1-associated RNAs were mapped to the human genome (GRCh37) and grouped to identify read clusters defining eIF4A1-bound regions ^46^. Binding was detected in 3082 and 9560 clusters per replicate (false discovery rate (FDR) = 0% and 0.42% respectively) with multiple clusters mapping to a single RNA (Fig. 5B). Following CMLD012824 treatment, there was a significant increase in the number of eIF4A1-bound clusters (161568 and 158503 per replicate; FDR 0.46% and 0.01% respectively), indicative of the magnitude of increased eIF4A1 association with RNA. We next characterized the nature of eIF4A1 binding clusters in these cells. The median length of the clusters in both DMSO- and CMLD012824-treated cells was 150 nucleotides (nts) or less (Fig. 5B). However, while these clusters formed the majority (85 ± 8%) of the eIF4A1-bound RNAs in DMSO-treated cells, they accounted for only half of the total clusters in CMLD012824-treated cells (51 ± 6%), with the remaining clusters being up to 1000 nts in length. A total of 826 and 1833 eIF4A1-bound RNAs per replicate was observed in DMSO-treated cells (165 consensus RNAs, FDR = 0%). Following CMLD012824 treatment, the number of eIF4A1-bound RNAs was substantially higher (13128 and 12593 transcripts per replicate), with 9269 consensus RNAs (FDR = 0.2%). The majority (∼80%) of eIF4A1 binding occurred at mRNAs of protein coding genes in both conditions, although a significant proportion (∼20%) also occupied long intergenic noncoding RNAs (lincRNAs) (Fig. S5D). We next investigated whether CMLD012824 treatment had any effect on the mRNA binding efficiencies of eIF4A1 by measuring the number of reads per transcript relative to the total RNA amounts^43^. Most mRNAs (∼90%, 1978/2212, *P* <0.1) became more strongly associated with eIF4A1 following CMLD012824 treatment relative to those in DMSO-treated cells (>1.5-fold change) (Fig. 5C). The CMLD012824-sensitive mRNAs were enriched for cell cycle and proliferation factors (G2/M transition, mitotic markers, mTORC1 signaling) as well as RNA regulation (RNA degradation, RNA binding), consistent with the functional enrichment of polypurine-rich genes correlated with amplified MYCN in primary NB tumors (Figs. S5E, S1E).

We next determined whether eIF4A1 binding was influenced by 5′ UTR sequence. Contrary to the well-known mechanism of action of rocaglates of disrupting translation initiation by clamping eIF4A1 to polypurine-rich 5′ UTRs ^30^, surprisingly only 27% of the CMLD012824-sensitive mRNAs were ranked as having polypurine-rich 5′ UTRs, suggesting a degree of stochastic binding (Fig. 5C). Alternatively, it also raised the possibility that polypurine content outside the 5′ UTR may account for the enhanced eIF4A1 binding.

As such, we first queried eIF4A1-mRNA interactions along the length of the transcripts and whether these were altered following ADR treatment. Under normal DMSO-treated conditions, as expected eIF4A1 binding was seen along the 5′ UTRs (∼15%), but greater numbers of clusters were found in coding sequences (CDSs; ∼50%) and 3′ UTRs (∼25%) (Fig. S5F). Only 1% of the eIF4A1-bound mRNAs showed binding throughout the entire lengths of the transcripts, and the 5′ UTR clusters were either unique (11 ± 3%; n = 2) or overlapped with the CDS (7.5 ± 1%, n = 2), but not the 3′ UTR (Fig. S5G). These patterns of binding were similar in CMLD012824-treated cells, suggesting that naïve eIF4A1 cluster distributions along mRNAs are largely retained upon ADR-mediated clamping, with a modest increase observed when binding included the 5′ UTRs or CDS together with the 3′ UTRs (Fig. S5F and S5G). Intriguingly, a large proportion of eIF4A1 binding was observed at the CDS (DMSO, 43±1%; ADR-treated, 39%, n = 2 each), in keeping with the recent observation that eIF4A1 may bind polypurine sequences in the CDS following RocA treatment ^47^. Aggregate read distribution analysis of eIF4A1 binding within the 5′ UTRs themselves showed that under normal conditions (DMSO treatment) binding gradually increased in the 3′ direction toward the start codons, whereas following CMLD012824 treatment eIF4A1 binding was more pronounced at the 5′ end of 5′ UTRs (Fig. 5D), further demonstrating the sustained clamping ability of the compound. These 5′ UTR-specific clusters in CMLD012824-treated cells were enriched for positive regulation of translation in response to stress, likely indicative of a compensatory response to translation inhibition (Figs. S5H).

Next, we determined whether ADRs mimicked the rocaglate predilection for polypurine-rich sequences by analyzing eIF4A1-bound mRNA sequences by *de novo* motif enrichment analysis ^48^. Interestingly, under normal conditions, a highly significant enrichment was observed for the [GAGA]_n_ and [AGG]_n_ polypurine motifs of eIF4A1-bound RNAs not only at the 5′ UTRs, but also along the entire length of the transcripts, pointing to the strong preference of eIF4A1 for polypurine sequences even in the absence of an inhibitor (Figs. 5E, S6A). On the other hand, in CMLD012824-treated cells, the significantly enriched motifs were [GAG]_n_ and [AAAA]_n_, suggesting that the ADR inhibitor selectively enhances eIF4A1 preference for polypurine sequences with a higher adenosine content (Fig. S6A). This was especially true for the subset of clusters that mapped to the 5′ UTRs (Fig. 5E). At the same time, the CDS- and 3′ UTR-specific binding clusters showed lesser enrichment for polypurine motifs, and, in addition, demonstrated higher enrichment for entirely novel C-containing motifs (Fig. 5E, S6B). Comparison of relative motif enrichment between DMSO- and ADR-824-treated datasets further supported the selective preference for adenosine content in the 5′ UTRs of the latter (Fig. S6C). Thus, the polypurine specificity of CMLD012824-mediated eIF4A1 clamping arises primarily at the 5′ UTRs and appears to be enriched for poly-(A) sequences. These findings suggest that eIF4A1 has an innate polypurine preference, may function at multiple locations along the mRNA and not only at the 5′ leaders, and while ADR treatment augments eIF4A1 binding and retains polypurine specificity in the 5′ UTR, it also exhibits variable specificity at other mRNA regions.

Given the striking sensitivity of MYCN-amplified NB cells to CMLD012824, we further analyzed eIF4A1 binding to the MYCN transcript. The eIF4A1 binding sites along the MYCN mRNA followed the overall binding pattern, with reproducible peaks appearing at the 5′ UTR, the coding region, and the stop codon in both DMSO- and CMLD012824-treated cells (Fig. 5F). However, the binding efficiency of eIF4A1 was significantly augmented with CMLD012824, with a ∼50-fold increase in binding peaks observed across the full length of the transcript in comparison to control cells (Fig. 5F). On the other hand, in contrast to MYCN and other polypurine-rich mRNAs, eIF4A1 binding along the 5′ UTRs of polypurine-poor transcripts such as PHOX2B was virtually absent even in CMLD012824-treated cells (Fig. 5G), indicating that ADRs also retain polypurine specificity.

We next sought to determine whether the observed changes in translation efficiency noted on ribosome profiling following ADR treatment could be attributed to the sustained clamping of eIF4A1 onto 5′ UTRs as determined by PAR-CLIP analysis. Of the 9269 eIF4A1-bound consensus RNAs in CMLD12824-treated cells, 30% (n = 2789) met statistical significance in the ribosome profiling results, of which 37% (n = 1037) and were translationally downregulated (>1.5-fold change). Interestingly, a number of eIF4A1-bound mRNAs were also translationally *up*regulated (n = 621, 22%, >1.5-fold change) (Fig. S6D). The eIF4A1-bound RNAs that corresponded to downregulated mRNAs were more highly enriched for polypurine-rich 5′ UTRs (29%; n = 297) compared to those associated with upregulated mRNAs (14%; n = 88) (Fig. S6D), consistent with observations in the total group of translationally regulated mRNAs (Fig. 4C). Comparison of differential translational efficiencies (Fig. 4C) and eIF4A1 clamping (Fig. 5C) between DMSO-treated and CMLD012824-treated cells revealed that 18% of translationally downregulated mRNAs exhibited increased eIF4A1 binding upon treatment (326/1841) (Fig. S6E). Here again, we also noted that a similar proportion of translationally *up*regulated mRNAs had increased eIF4A1 binding (16%, n = 234/1451) (Fig. S6E). The relatively low numbers of significant mRNAs in this integrative analysis are likely due to the caveat of comparing ribo-seq data from long-established Kelly cells and PAR-CLIP from COG-N-415x PDX cells, although both cell types express amplified MYCN. Taken together, these findings delineate the features of CMLD012824-sensitive and insensitive mRNAs and suggest parameters for predicting whether select mRNAs are inhibited, remain unaffected or even achieve upregulation.

### CMLD012824 clamps EIF4A1 onto select polypurine-rich cellular mRNAs in a 5′ UTR-dependent manner

Although eIF4A1 exhibited surprisingly promiscuous mRNA clamping beyond the 5′ UTRs in NB cells, which was augmented along the full lengths of the mRNAs following CMLD012824 treatment, we sought to determine whether the 5′ UTR region alone is sufficient to confer the observed effect on translation. We therefore investigated the direct effects of CMLD012824-mediated eIF4A1 binding to endogenously expressed 5′ UTRs in NB cells. First, to test the functional role of the 5′ UTR in translation of the MYCN protein, we overexpressed a human MYCN cDNA construct lacking the 5′ UTR in MYCN-nonamplified NB cells (Fig. 6A). Without the endogenous MYCN 5′ UTR, CMLD012824 had no activity against MYCN protein levels, indicating that this region was necessary for the translation-inhibition effect of the ADR (Fig. 6A). Next, we questioned whether an endogenous polypurine-rich 5′ UTR would be sufficient for CMLD012824-mediated translation inhibition through an *in vitro* translation assay. In agreement with the ribosome profiling results (Fig. 4C), CMLD012824 inhibited the translation of a luciferase reporter downstream of not only the MYCN 5′ UTR but also of other polypurine-rich 5′ leaders such as JUN and CCND1 (Fig. 6B). By contrast, translation from polypurine-poor controls (CKS2, XRN2) was not affected by CMLD012824, as was an eIF4A scanning-independent control, the hepatitis C viral internal ribosome entry site RNA (HCV IRES) ^49^ (Fig. 6B).

**Figure 6.**
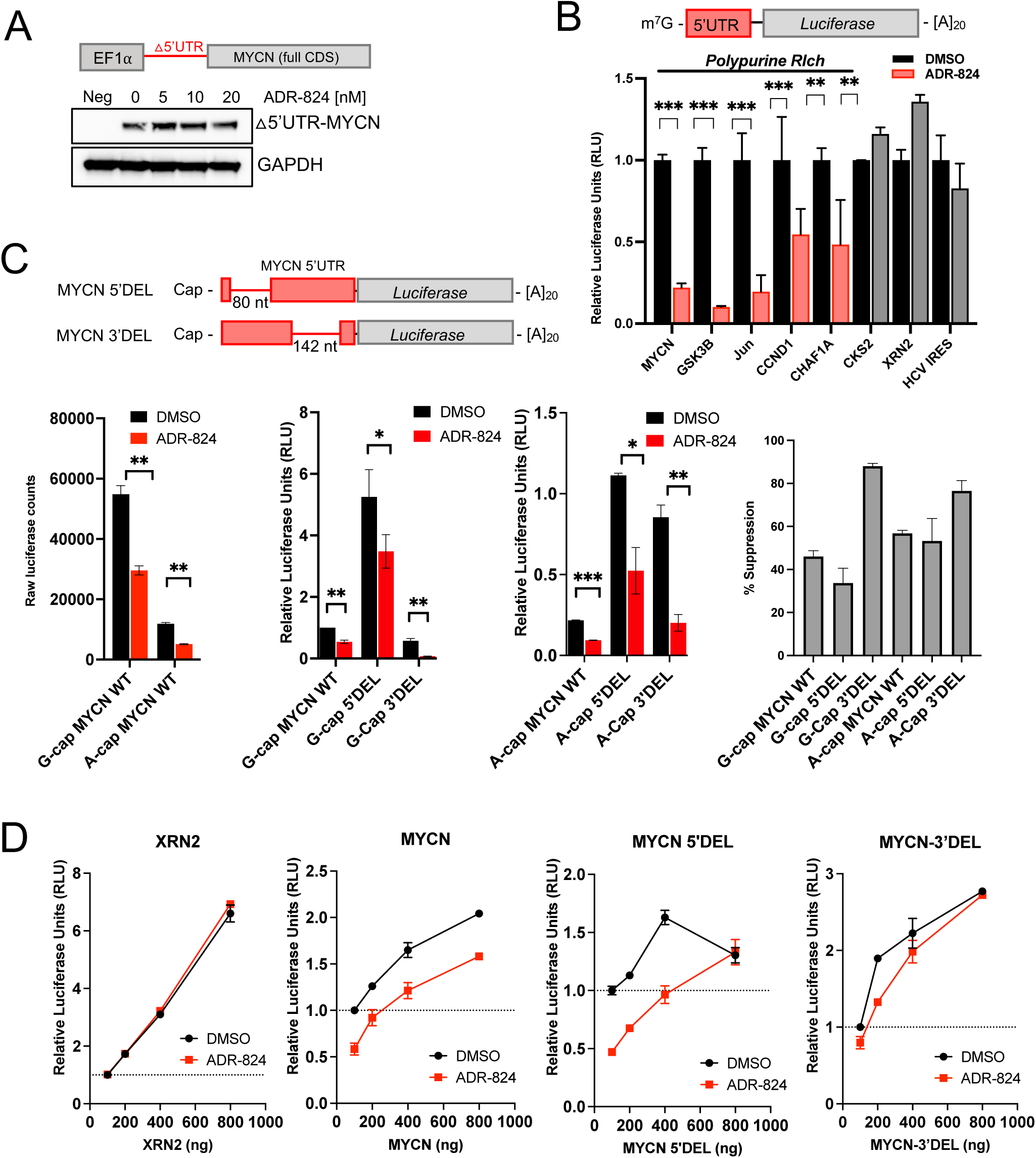
ADR-824 clamps eIF4A1 onto select polypurine-rich cellular mRNAs in a 5′ UTR- dependent and cap-independent manner. **A.** WB analysis of exogenously expressed 5′ UTR-depleted MYCN in MYCN-non-expressing SK-N-AS NB cells, treated for 1 h with the indicated doses of ADR-824. GAPDH serves as the loading control. Schematic above depicts deletion of the endogenous 5′ UTR, leaving a construct of full-length MYCN CDS under a mammalian promoter. B. *Renilla* luciferase activity from *in vitro* translation of endogenous 5′ UTR sequences inserted upstream of *Renilla* in the presence of DMSO or ADR-824 (25 nM). Signal is normalized to internal globin-*Firefly* luciferase control. CKS2 and XRN2 serve as polypurine-poor controls; HCV IRES RNA serves as an eIF4A-independent control. Data represent mean ± S.D., n = 3 replicates. **p value < 0.001, ***p value < 0.0001, Student’s t-test. **C.** Top: schematic representation of the wild type MYCN 5′ UTR sequence, 5′ deletion mutant (MYCN 5′ DEL), and 3′ deletion mutant (MYCN 3′ DEL) inserted upstream of *Renilla* luciferase. Bottom left (three panels): *in vitro* translation of indicated RNAs generated through canonical m^7^G-cap (G-cap) or nonfunctional ApppG analog (A-cap) in the presence of DMSO or ADR-824. Bottom right: percent suppression of translation measured by luciferase activity. Data represent mean ± S.D., n = 2 replicates, representative of 3 independent experiments. *p value < 0.01, **p value < 0.001, ***p value < 0.0001, Student’s t-test. D. *Renilla* luciferase activity from *in vitro* translation of indicated RNAs at the indicated concentrations in the presence of 200 ng per reaction of globin-*Firefly* RNA with DMSO or ADR-824 (25 nM). Data represent mean ± S.D., n = 3 replicates.

Although canonical cap-dependent translation is by far the most prevalent mechanism of translation initiation in mammalian cells, non-canonical modes such as internal ribosome entry sites (IRES) can be utilized by cancer cells exposed to hypoxia or cytotoxic stress ^50^. Indeed, translation of several oncogenic transcription factors, including MYCN, has been shown to be initiated via IRES elements within their 5′ UTRs ^37^. We therefore determined whether the observed effect of CMLD012824 was primarily through disruption of cap-dependent translation initiation or whether IRES-driven activity was also affected. We analyzed the effects on luciferase reporter activity from wild type (WT) and IRES-mutant MYCN 5′ UTRs in the presence of a canonical m^7^G-cap or a non-functional A-cap, of which the latter is only translatable through IRES activity, under untreated or CMLD012824-treated conditions. We generated two different cDNA constructs with deletions of either 80 nucleotides (nts) at the 5′-end (MYCN 5′ DEL) or 142 nts at the 3′-end (MYCN 3′ DEL) of the 5′ UTR, both of which have been suggested to confer IRES activity in bicistronic assays ^37^. We then generated *in vitro* transcribed MYCN WT and deletion-mutant mRNAs containing a non-functional 5′ ApppG cap analog (A-cap) in place of the canonical functional m^7^GTP cap structure ^51^. The A-capped MYCN WT 5′ UTR retained ∼20% luciferase activity relative to the m^7^G-capped UTR (Fig. 6C), indicative of IRES activity, which was inhibited by a further 10% with CMLD012824 (Fig. 6C). Compared to MYCN WT, removal of the 5′ IRES segment (MYCN 5′ DEL) significantly de-repressed translation from the 5′ UTR relative to MYCN WT of m^7^G-capped and A-capped mRNAs, suggesting that this region serves an inhibitory function (Fig. 6C). Removal of the 3′ segment (MYCN 3′ DEL) resulted in decreased translation of the m^7^G-capped RNA. On the other hand, this segment showed increased translation of the A-capped mRNA (Fig. 6C), suggesting that IRES activity is retained^37^. Treatment with CMLD012824 inhibited the translation of not only the wild type MYCN 5′ UTR but also both the 5′ and 3′ deletion mutants (Fig. 6C) suggesting that the entire MYCN 5′ UTR contains ADR-sensitive sequences. Importantly, CMLD012824 inhibited the translation of all the A-capped mRNAs, indicating that ADR-mediated inhibition also extends to non-canonical translation events (Fig. 6C).

To prove whether the relative upregulation of mRNAs following translation inhibition as observed in the analysis of our ribo-seq and PAR-CLIP analyses could be explained by the length and polypurine composition of the 5′ UTR, we performed *in vitro* translation competition assays against the short polypurine-poor globin 5′ UTR in the presence of CMLD012824 or DMSO control. The short, polypurine-poor 5′ UTR of the XRN2 gene was able to compete efficiently against the globin 5′ UTR under both DMSO- and CMLD012824-treated conditions (Fig. 6D). By contrast, the MYCN 5′ UTR was consistently inhibited by CMLD012824 and could not be overcome even at higher RNA concentrations (Fig. 6D). Competition against the MYCN 5′ DEL and MYCN 3′ DEL mRNAs, however, resulted in a near-total rescue of the effect of CMLD012824, with the deletion mutants competing against the globin 5′ UTR at higher concentrations (Fig. 6D). Rescue with the MYCN 3′ DEL RNA was more effective than with MYCN 5′ DEL, consistent with the removal of a larger number of polypurine nucleotides at the 3′ end of the 5′ UTR in comparison to the 5′ end (Fig. 6D). These results are in line with our ribo-seq and PAR-CLIP findings that suggest a dynamic aspect to ADR-mediated inhibition, where 5′ UTR content as well as competition between variable lengths and nucleotide compositions determine the outcome.

### CMLD012824 slows tumor growth in vivo and improves survival

To investigate whether ADR inhibitors could be a viable therapeutic option in NB, we tested the effects of CMLD012824 in several mouse models. As CMLD012824 has not previously been tested *in vivo* in enantiomerically pure form, we first established the maximum tolerated dose in non-tumor bearing C57BL/6J mice (Fig. S7A). We determined that a 0.1 mg/kg daily dose was well tolerated and was sufficient to induce a decrease in target protein levels (CCND1, CCNE1, CDK4) in liver tissue from treated animals, while eIF4A1 levels were unchanged, as observed in our *in vitro* studies (Figs. S7B, 1G, 2D). We next tested the compound in murine xenograft models of NB-9464 cells derived from the TH-MYCN transgenic mouse model of MYCN-driven NB ^52,53^.

Cells were inoculated subcutaneously into the flanks of syngeneic C57BL/6J mice and upon tumor formation, the animals were treated with vehicle and CMLD012824 (two doses, 0.1 and 0.2 mg/kg) three times a week by intraperitoneal injection for 30 days (Fig. 7A). While animals treated with both doses of CMLD012824 exhibited no toxicities, a reduction in tumor burden was observed for those treated with the higher dose although the study was not adequately powered to establish significance (Fig. 7A). Nevertheless, we still observed loss of MYCN protein, as well as decreased levels of another polypurine-rich 5′UTR target (DDX1), at both doses, while the polypurine-poor control (CKS2) was upregulated in tumors from CMLD01284-treated mice (Fig. S7C). Finally, we tested the *in vivo* effects of CMLD012824 in a PDX model of MYCN-amplified NB (COG-N-415x) generated in nude mice. Vehicle or 0.2 mg/kg CMLD012824 was administered three times a week by intraperitoneal injection until endpoint tumor volume was reached (>1000 mm^3^) or completion of the study (50 days). A significant decrease in tumor size was observed in mice treated with CMLD012824 (Fig. 7B), with an improvement in overall survival (Fig. 7C). Immunohistochemistry (IHC) analysis confirmed decreased tumor proliferation and increased apoptosis in response to CMLD012824 treatment, as measured by Ki67 and cleaved caspase 3 staining respectively (Fig. 7D, S7D) as well as a clear downregulation of MYCN protein levels (Fig. 7D, S7D). Western blot analysis confirmed loss of MYCN and DDX1, as well as JUN, a polypurine-rich 5′ UTR target (Fig. 1G), while polypurine-poor controls (CKS2, XRN2) and initiation factors (eIF4A1, eIF4E) remained unaffected (Fig. 7E, S7E). These studies together demonstrate that the ADR derivative CMLD012824 causes inhibition of tumor growth in MYCN-driven NB models with tolerable toxicity.

**Figure 7.**
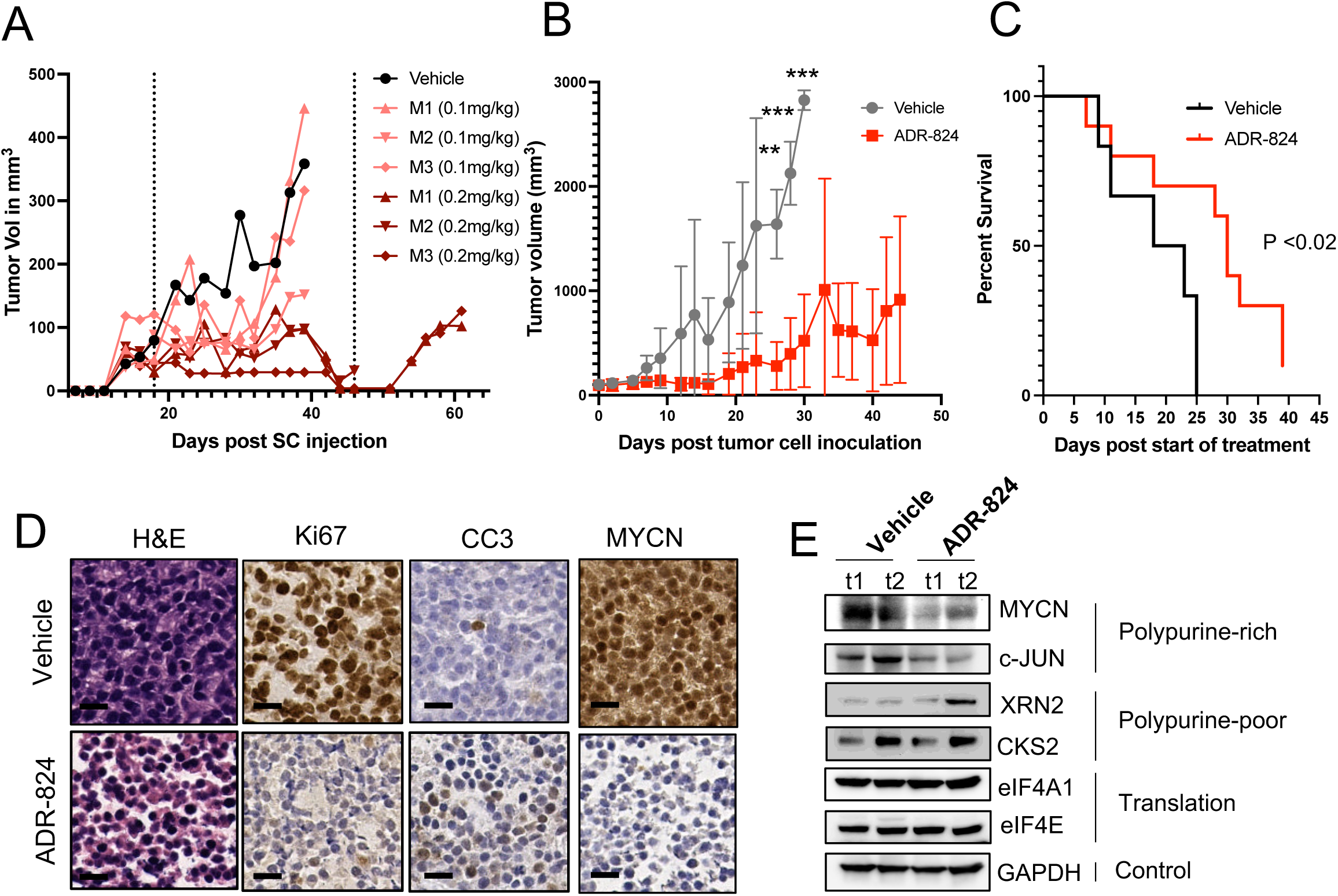
ADR-824 inhibits tumor growth and improves survival in NB. **A.** Tumor volumes of NB-9464 xenograft tumors in C57BL/6J mice (n=10) treated three times a week (Mondays, Wednesdays, Fridays) with the indicated doses of ADR-824. Dashed lines indicate beginning and end of treatment. Each curve corresponds to a separate animal, (vehicle vs 0.1 mg/kg, *P* < 0.25; vs 0.2 mg/kg, *P* < 0.006, Welch’s test). **B.** Tumor volumes of patient-derived MYCN-amplified (COG-N-415x) xenografts treated three times a week (Mondays, Wednesdays, Fridays) with vehicle (n=7) or ADR-824 (n=10). Data represent mean ± S.D. Statistically significant differences between treatment groups were observed on days 21, 23, and 25 (*p < 0.01, **p < 0.001, ***p < 0.0001, Student’s t-test) after which no vehicle treated animals survived. **C.** Kaplan-Meier analysis of COG-N-415x PDX-bearing mice (B) (*P* < 0.02, Mantel-Cox t test). **D**. Representative images of hematoxylin and eosin (H&E) and immunohistochemistry analyses (IHC) of the indicated tumor markers [Ki67 (proliferation), CC3, cleaved caspase 3 (apoptosis)] in vehicle (top) and ADR-824-treated (bottom) mice. Scale bar, 100 µm. **E.** WB analysis of the indicated polypurine-rich, -poor, translation factor and control proteins in COG-N-415x PDX tumors in (D) and (S4D). t# – tumor designation in group.

## Discussion

Direct targeting of amplified MYCN has proven to be challenging in neuroblastoma. Here, we demonstrate that targeting the complex 5′ UTR regulatory elements of MYCN at the level of translation using novel ADRs could offer an alternative route for disrupting the aberrant proliferative activities of this oncogene. Moreover, ADR-mediated translation inhibition provides an avenue for inhibiting a functionally important subset of cellular mRNAs – those critical to malignant proliferation - many of which, like MYCN, are enriched in ADR-sensitive polypurine-rich motifs. Such selective suppression of protein synthesis is enabled by the transcriptional landscape and unique gene dependencies of MYCN-amplified NB, where the extreme reliance of deregulated MYCN on protein synthesis creates a vulnerability that when targeted, leads to selective cytotoxicity while sparing normal tissue.

The synthetic rocaglate CR-1-31-B has previously been shown to be cytotoxic in two NB cell lines^31^, however, its impact on amplified MYCN and the mechanisms underlying such a response are unclear. We show that the novel ADR CMLD012824 is not only highly potent against multiple MYCN-amplified NB cell lines, but also, importantly, demonstrate its tolerability and efficacy in animal models, thereby providing pre-clinical validation for further development of this class of inhibitors. In addition, we interrogate the effects of this rocaglate derivative on MYCN-driven protein synthesis and demonstrate that the long and polypurine-rich 5′ UTR containing a predicted complex secondary structure that requires unwinding by eIF4A1^18^ renders MYCN an ideal ADR target. MYCN as well as other essential transcription factors with mRNAs that possess polypurine-rich 5′ UTRs, would be expected to be sensitive to ADR-mediated inhibition, leading to a feed-forward deregulatory loop as the translation of these drivers of proliferation is blocked and their effects on transcription are lost. Indeed, this is precisely what we observe through ribosome profiling analysis which revealed ADR-driven decreases in translation efficiencies of mRNAs corresponding not only to MYCN, but also to other DNA-binding regulatory factors, as well as drivers of major proliferative signaling pathways in NB. Additionally, super-enhancer associated^5^ and MYCN target^44^ mRNAs, which are central to driving oncogenic transcription, were sensitized according to the polypurine content of their 5′ UTRs, suggesting that polypurine ranking of key oncogenic targets can be predictive of the downstream magnitude of ADR-mediated inhibition. This predictive power is exemplified by the comparison of MYCN and c-MYC mRNAs, where the higher polypurine content of the MYCN 5′ UTR results in stronger ADR-mediated inhibition and greater loss of the protein. The lack of effect on c-MYC protein levels was also recently reported in pancreatic ductal adenocarcinoma models that were sensitive to CR-1-31-B^24^. Moreover, the significant correlation of amplified MYCN with eIF4F complex expression in primary NBs (Figs. 1A-D) signals the selective advantage ADRs would have in cells in which MYCN is the driver. The concomitant decreases in corresponding protein levels of the genes that regulate the proliferative network reasonably accounts for the cellular cytotoxicity and effects on tumor burden in animal models.

We also describe the characteristics of both ADR-sensitive and -insensitive mRNAs, highlighting the importance of interrogating the mRNA sequence, length, binding motifs and additional features such as cap dependence to delineate the target preferences of rocaglate analogs. The CMLD012824-insensitive mRNAs were characterized by short, polypurine-poor 5′ UTRs, that are likely to be less dependent on eIF4A1 activity^54,55^. This finding further illustrates the variable specificity of rocaglate compounds, as prior studies have identified G-quadruplexes^17^, low GC content (silvestrol)^26^, high GC content (hippuristanol) ^20^, short 5′ UTRs^24^ or cap-dependent polypurine targeting (CR-1-31-B)^30^ as major determinants of inhibitory activity. While analysis of the ADR effect on protein translation confirmed the expected decrease in translation efficiencies of polypurine-rich mRNAs and demonstrated a stronger inhibitory effect on longer 5′ UTRs, PAR-CLIP analysis generated a comprehensive map of the distribution of naïve and ADR-bound eIF4A1 protein on cellular mRNAs. Recent studies have shown that eIF4A1 interacts not only with the eIF4F complex but is capable also of loading independently onto mRNAs ^15,36^. In agreement, we observed that naïve eIF4A1 associates promiscuously with cellular mRNAs outside of 5′ UTR regions, suggesting that eIF4A1 may exhibit stochastic associations with mRNAs independent of the cap-dependent eIF4F complex. Importantly, we find that eIF4A1 exhibits preferential polypurine binding even in the absence of the inhibitor, implying that eIF4A1 spends more time sampling polypurine-rich rather than pyrimidine-rich sequences under normal conditions. On the other hand, ADR-mediated eIF4A1 binding exhibited more complexity. While binding was greatly augmented by the ADR along the full lengths of mRNAs, increased clamping was observed at the 5′ termini of the 5′ UTRs, particularly of those corresponding to proliferative mRNAs such as MYCN, CCND1 and JUN, and suggestive of blocks to pre-initiation complex scanning and subsequent translation initiation. We also noted a surprising preference of eIF4A1 toward poly-adenosine sequence motifs in the 5′ UTRs of bound mRNAs in ADR-treated cells, while CT-containing motifs were enriched in the CDS or 3′ UTRs. One explanation for this observation is that the high-affinity purely polypurine sites have become saturated with eIF4A1 and the excess eIF4A1 is being bound to lower affinity (CT-containing) sequences. Alternatively, other factors such as RNA binding proteins (RBPs) could contribute to the differing sequence specificity of ADRs between mRNA regions. Overall, mRNAs that exhibited increased eIF4A1 binding upon ADR treatment were noted to be more likely to be translationally downregulated. However, as previously observed ^33^, ADR-mediated clamping of eIF4A1 did not fully correlate with the loss of translation efficiency, likely due in part to the different models analyzed using ribosome profiling and PAR-CLIP. Alternatively, the eIF4A1 clamping observed in the 3′ UTR regions may not contribute to decreases in translation, but rather may interfere with microRNA mediated inhibition, such as in the case of MYCN^56^. As has been suggested^47^, sustained clamping of eIF4A1 in the CDS may also modulate translation elongation, thereby accounting for the lack of correlation between 5′ UTR binding and translation efficiency. The significant degree of clamping we observed in the CDS and 3′ UTRs may also result in sequestration of eIF4A1, causing the cytotoxic effect of the ADR to be compounded by reducing the amount of free eIF4A1 that is available for translation of mRNAs without polypurine-rich motifs ^30,57^. Curiously, mRNAs which were translationally upregulated in both MYCN-amplified and nonamplified ADR-treated cells were enriched for translation elongation and termination factors, as well as mitochondrial translation machinery, pointing to potential compensatory responses to the additional translation defects (Fig. S3C). These results, backed by the *in vitro* demonstration of the requirement of the 5′ UTR for translation inhibition, suggest that inhibition of eIF4A1 at the 5′ UTRs of target mRNAs, and not the CDS or 3′ UTRs, primarily confers ADR-inhibitory activity. Meanwhile, the association of eIF4A1 with the CDS, 3′ UTRs, as well as other types of RNAs, particularly those augmented by the ADR, suggests the possibility of complex secondary effects that warrant further study.

Our studies also revealed an unexpected aspect of CMLD012824-mediated inhibition - increased translational efficiency of a subset of mRNAs, even in the presence of increased eIF4A1 clamping. While the identification and classification of sensitive mRNAs (long, polypurine-rich 5′ UTRs without cap dependence) is critical to identifying the direct targets of CMLD012824, characterization of insensitive mRNAs (short, polypurine-poor 5′ UTRs) is vital to deciphering the global effects of ADRs *in vivo*. Previous models of rocaglate-mediated target inhibition have suggested a multi-modal effect in which the dominant-negative clamping of the rocaglate on target mRNAs is coupled with a bystander effect where off-target mRNAs are inhibited due to a decrease in available translation machinery^30,36^. Our results expand on this model of ADR-mediated translation inhibition by demonstrating that short polypurine-poor mRNAs not only escape inhibition but also outcompete longer, more complex transcripts to become upregulated. We postulate that the competition effect arises from the limiting amount of available translational machinery, which is not sequestered in a dominant-negative manner by the ADR-mediated clamping of sensitive mRNAs. Transcripts that have short, unstructured 5′ UTRs can more effectively compete for the remaining translation initiation complexes and are consequently translationally upregulated. The competition of variable 5′ UTR compositions is revealed by the selective ADR inhibition and is suggestive of the competition between cellular mRNAs for translation machinery under normal conditions. Importantly, the translational capability of endogenous 5′ UTR regions can be improved by the removal of polypurine content, as we demonstrate with the MYCN 5′ UTR. The increase in translational efficiency of short, polypurine-poor mRNAs can be predicted using our polypurine ranking model, analogous to that of ADR-mediated downregulation based on polypurine-rich mRNA composition. This concept can thus be applied to anticipating sensitivities to ADR treatment in various cancers, which may depend on oncogenic factors that are selectively downregulated, such as MYCN, or may escape ADR-mediated inhibition (*e.g.* PHOX2B) due to the 5′ UTR composition of their respective mRNAs. Further studies will reveal the impact on other DEAD-box helicases (*e.g.* eIF4A2) which may be ADR targets and whose inhibition may result in additional cytotoxic effects, as well as potential compensation and resistance mechanisms.

In summary, our study describes a novel strategy for overcoming the oncogenic effects of amplified MYCN, whose direct targeting has thus far been unsuccessful. The specific dependence of MYCN-amplified NB cells on increased protein synthesis together with the unique mRNA selectivity of ADRs results in the preferential targeting of NB cells *versus* normal tissues. Our results provide preclinical evidence for using ADRs in the treatment of MYCN-amplified NB and describe how this strategy may be implemented in other transcriptionally driven cancers.

## Acknowledgements

The results shown here are in part based on data curated by the R2: Genomics Analysis and Visualization Platform: http://r2.amc.nl/. We thank members of the George lab for helpful discussions, Sanjukta Das for initial analysis of ChIP-seq data and Akiko Shimamura for the use of sucrose gradient fractionation equipment. We thank the late Jerry Pelletier for advice and reagents. This work was supported by a Friends for Life Neuroblastoma Foundation grant (R.E.G.), NIH grants R35GM118173, U01TR002625R21 (J.A.P., Jr. and L.E.B), R21CA267621 (R.E.G.) and an Alex’s Lemonade Stand Foundation Innovation grant 21-24757 (R.E.G).

## Author contributions

M.V. and R.E.G. conceived the study and designed the experiments. L.E.B and J. A. P Jr generated compounds and provided valuable feedback. M.V. performed the molecular, cellular and biochemical studies. M.V. performed the computational analysis with inputs from R.D., U.B and R.E.G. B.S. and M.V. performed the animal experiments. A.K. contributed to the polysome profiling. M.V. and R.E.G. wrote the manuscript with input from all authors.

## Supplementary Figure Legends

**Figure S1. Translation initiation machinery genes are correlated with MYCN but not c-MYC overexpression. A**. Hierarchical clustering of translation initiation factor gene expression in primary NB tumors (n=498, derived from GSE62564) grouped by annotated MYCN amplification status. Z-score mean ± S.D. Bar above heatmap represents corresponding MYCN expression level in log reads per million (log2 RPM). **B.** Violin plots showing expression of the indicated initiation factors in tumors with amplified and nonamplified MYCN, as depicted in A. Box plots within the violin plots defined by center lines (medians), box limits (the interquartile range between 25th and 75th percentiles), whiskers (minima and maxima). Significance determined by Student’s t-test. **C.** Correlogram of c-MYC and translation initiation factor gene expression in MYCN-nonamplified NBs (n = 401, GSE62564). Circles represent Spearman’s rank correlation coefficients, P <0.01. Color code represents positive correlations in blue, negative correlations in orange-red. **D.** Dot plot showing highly variable genes (HVG) identified from gene expression data in primary NB tumors (n=498, GSE62564). Variance was determined by arraying and binning all genes by expression level and calculating the variance coefficient for each group, which was then converted into a z-score. Significant HVGs with a z-score > 0.1 are depicted in gold. **E.** Volcano plot showing changes in expression of highly variable genes (HVGs) in tumors with lowest and highest MYCN expression levels (n=30 each, GSE62564). Y-axis shows significance of variance in log P-value, with horizontal line representing cutoff of *P* < 0.01. X-axis shows log2 fold change. Upregulated genes (>2-fold change) are shown in red, downregulated genes in blue, Student’s t-test, *P* < 0.01. **F.** Fold change distributions of highly variable genes (HVGs) in tumors from lowest (black, bins 1 and 2) to highest (red, bins 9 and 10) MYCN expression levels (n=30 each, GSE62564) (Student’s t-test, P < 0.01). X-axis shows the magnitude and Y-axis shows the frequency of fold change (FC) in expression. **G**. Contour plot showing the polypurine rank distribution of highly variable genes (HVGs). **H.** Polypurine rank distribution of the highly variable upregulated genes from lowest (black, bins 1 and 2) to highest (red, bins 7 and 8) fold change. (Student’s t-test, P < 0.01). **I**. Functional enrichment analysis of the top 25% polypurine-rich genes ranked by MYCN expression in NB primary tumors (n=498). FDR <0.1.

**Figure S2. CMLD012824 leads to differential cytotoxicity in NB.**

**A.** Cell viability of MYCN amplified (red) and non-amplified (blue) human PDX-derived NB cells, treated with varying concentrations of CMLD012824 (ADR-824) for 72 h. Percent cell viability relative to DMSO is shown. Data represent mean ± S.D., n = 3 replicates. **B.** Annexin V (left) and membrane integrity (right) analysis at the indicated times following treatment with the indicated doses of ADR-824 in MYCN-amplified (CHP-134) cells. Data represent mean ± S.D., n = 3 replicates. **B.** Quantification of propidium iodide incorporation in the indicated NB cells 72 h post treatment with ADR-824 at the indicated doses. Data represent ± S.D., between n = 3 biological replicates.

**Figure S3. ADR-824-mediated inhibition targets unique subsets of long, poly-purine-rich mRNAs in MYCN-amplified versus nonamplified NB cells.**

**A.** Scatter plot of total vs. ribosome associated mRNA changes in ADR-824-treated MYCN-nonamplified (SK-N-AS) NB cells (10 nM, 1 h), with axes showing log2 fold change as determined by Anota2seq (P <0.1) (See Methods). Dotted black lines indicate cut-offs of 1.5-fold change in total mRNA (x-axis) and ribosome occupancy (y-axis) of ADR-824-treated biological replicates (n=3) compared to vehicle control (n=3). Data points are color coded according to ribosome occupancy, with corresponding total numbers shown on the plot (Upregulated shown in gold, downregulated shown in blue). **B.** Venn diagram representing the overlap in translationally regulated MYCN-amplified (Kelly) and nonamplified (SK-N-AS) gene sets. Down: >1.5-fold decrease, Up: >1.5-fold increase (*P* <0.1). **C.** Functional enrichment of common downregulated (top, blue) and upregulated (bottom, red) processes in MYCN-amplified and non-amplified NB cells identified by ribosome profiling analysis [Down: >1.5-fold decrease, Up: >1.5-fold increase (*P* <0.1)]. **D.** Functional enrichment of unique downregulated processes in non-amplified (SK-N-AS) cells [Down: >1.5-fold decrease (*P* <0.1)]. Axis depicts log of inverse P-value in **C, D**. **E.** WB analysis of representative unique upregulated mRNAs in MYCN-amplified (Kelly) cells, with GAPDH as control. **F.** Volcano plot of translationally regulated mRNAs in ADR-824-treated MYCN-nonamplified (SK-N-AS) NB cells, showing translational efficiency changes as determined by Anota2seq (>1.5-fold change, *P* <0.1). Polypurine-rich (top 25%) mRNAs are highlighted in red, with percentages indicated on the plot. **G.** Motif enrichment analysis showing top motifs in the downregulated subset, trained against a background list of unregulated transcripts. **H**. Ranking of super-enhancer associated genes in MYCN-amplified (Kelly) NB cells based on histone 3 lysine 27 acetylation (H3K27ac) signal, plotted in order of increasing super-enhancer rank (MYCN=1, highest). Translational efficiency changes and polypurine ranking were calculated as in (4C). Upregulated mRNAs are shown in gold, downregulated in blue, >1.5-fold change Polypurine-rich (top 25%) mRNAs are highlighted in red. Right: quantification of polypurine-rich (top 25%) and polypurine-poor (bottom 25%) in the downregulated mRNA subset.

**Figure S4. ADR-824-mediated inhibition alters ribosome occupancy of poly-purine-rich mRNAs in MYCN-amplified NB cells. A.** Contour plot showing 5′ UTR length distribution of translationally regulated transcripts in (gold: upregulated, blue: downregulated) in ADR-824-treated MYCN-nonamplified (SK-N-AS) NB cells. *P* <2.2e-16 (up- vs. down-regulated mRNAs, >2-fold change, Student’s t-test). **B**. Contour plot of polypurine rank distribution of translationally regulated transcripts in nonamplified (SK-N-AS) NB cells. *P* <2.2e-16; Student’s t-test. **C, D.** Scatter plot of polypurine content (**C**) and GC content (**D**) by 5′ UTR length in upregulated (gold) versus downregulated (blue) mRNAs in nonamplified cells. Loess regression analysis is shown in corresponding colors, shaded regions represent 95% confidence intervals. **E.** A_260_ absorbance signal of RNA in sucrose gradient sedimentation fractions. Kelly cells were treated for 1 h with vehicle or ADR-824 (10 nM), lysed and fractionated on 10-50% sucrose gradients by ultracentrifugation. The positions of 80S, light (L1-3) and heavy (H1-2) polysomes are indicated on the plot. Results show one representative experiment. **F.** RT-qPCR analysis of MYCN and HERC2 polypurine-poor control mRNA distribution in polysome fractions after vehicle or ADR-824 treatment in MYCN-amplified (Kelly) NB cells. Light polysomes: L1-3; heavy polysomes: H1-2. Signal was calculated by 2^-ΔΔCt method, normalized to total RNA in gradient. Data represent mean ± S.D., n = 3 replicates.

**Figure S5. eIF4A1 binds along the full length of mRNAs and other classes of RNAs. A.** WB analysis of eIF4A1 and the indicated proteins (with GAPDH used as a control) in immunoprecipitates of eIF4A1 or IgG from DMSO- and ADR-824-treated MYCN-amplified (Kelly) cell lysates. **B.** RT-qPCR analysis of 5′ UTR polypurine-rich and -poor mRNAs bound to eIF4A1 immunoprecipitated from MYCN-nonamplified (SK-N-AS) cell lysates following treatment with DMSO or ADR-824 (10 nM x 1 h). Data represent mean ± S.D., n = 3 replicates. **p value < 0.001, ***p value < 0.0001, Student’s t-test. **C**. WB analysis of immunoprecipitation (IP) of eIF4A1 after PAR-CLIP, with GAPDH serving as loading and crosslinking (negative) control. **D.** Distribution of eIF4A1-binding clusters in DMSO- and ADR-824-treated cells that map to the indicated classes of RNAs. **E.** Functional enrichment analysis of eIF4A1-bound clusters from ADR-824-treated cells. Combined score indicates P-value derived from Fisher’s exact test in Enrichr (*P* < 0.001 cutoff). **F**. Distribution of eIF4A1-binding clusters that map to the indicated mRNA regions. **G**. Distribution of eIF4A1-binding clusters to the indicated regions of mRNAs in ADR-824-treated cells. Average percentages out of the total unique mRNAs per replicate (DMSO, n = 826, 1833; ADR, n= 13128, 12593) indicated on right. **H**. Functional enrichment analysis of eIF4A1-binding clusters from ADR-824-treated cells that map to the 5′ UTRs. Combined score indicates adjusted p value derived from Fisher’s exact test in Enrichr (*P* < 0.001 cutoff).

**Figure S6. eIF4A1 binds to polypurine motifs along the full length of mRNAs.**

**A.** Top motifs identified in eIF4A1-binding clusters in DMSO- and ADR-824-treated cells. E-values adjusted to motif frequency. **B.** Additional top discovered motifs from ADR-824-treated eIF4A1- binding clusters that map to the indicated mRNA regions. **C.** Top motifs identified in eIF4A1 clusters that map to the indicated mRNA regions in ADR-824-treated cells, trained against the DMSO-treated background. E-values adjusted to motif frequency are shown. **D.** Left, Volcano plot of translationally regulated mRNAs (identified through ribo-seq) that correspond to eIF4A1-bound RNAs (identified through PAR-CLIP) in ADR-824-treated cells. The x-axis shows the ribosome profiling (RP) translational efficiency and y-axis, statistical significance, as determined by Anota2seq (>1.5-fold change, *P* <0.1). Polypurine-rich (top 25%) mRNAs are highlighted in red. Right: Quantification of polypurine-rich mRNAs with >2-fold change in translational efficiency, in down- and upregulated ribo-seq subsets. **E.** Scatter plot of ribosome-associated mRNA changes (x-axis) versus eIF4A1 PAR-CLIP binding changes (y-axis) from ADR-824-treated cells, with axes showing log2 fold change as determined by Anota2seq (*P* <0.1). Data points are colored according to changes in ribosome occupancy and eIF4A1 binding (>1.5-fold change in both) with corresponding numbers shown on the plot. Dotted black lines indicate cut-offs for ADR-824-treated compared to DMSO-treated biological replicates (1.5-fold change, n=2 each).

**Figure S7. Establishment of maximum tolerated dose of ADR-824 in murine models.**

**A.** Serial weights of C57BL/6J non-tumor-bearing mice treated daily for 5 days with the indicated doses of ADR-824 or vehicle. Weights are shown in grams ± S.D. n = 2 per ADR-824 treated, n=1 for vehicle-treated. Weight of zero indicates animal death. **B.** WB analysis of proliferation and cell cycle proteins (CCND1, CCNE1, CDK4) and translation initiation regulatory proteins (eIF4A1, eIF4E, 4E-BP1) in liver tissue of vehicle and ADR-824 (0.1mg/kg) mice in (A). GAPDH serves as a loading control. **C.** WB analysis of polypurine-rich (MYCN, DDX1), -poor (CKS2), and GAPDH control proteins in NB-9464 PDX tumors in Fig 7A. t# – tumor designation in group. **D.** H&E and IHC analyses of additional representative vehicle (top) and ADR-824-treated (bottom) COGN-415x-derived tumors. Scale bar represents 100 µm. **E.** WB analysis of polypurine-rich (MYCN, DDX1), -poor (CSK2, XRN2), and control (eIF4E, GAPDH) proteins in COG-N-415x PDX tumors in Fig 7A. t# – tumor designation in group.

## Materials and Methods

### Cell Culture

Human neuroblastoma (NB) cells (Kelly, IMR-32, CHP-134, NGP, GIMEN, LAN-5, SK-N-SH, SH-SY5Y, COG323, COG327, COG346, COG415, COG476, COG504, COG514) were obtained from the Children’s Oncology Group cell line bank and genotyped at the Dana-Farber Cancer Institute (DFCI) Core Facility. NB cells were grown in RPMI (Invitrogen) supplemented with 10% FBS and 1% penicillin/streptomycin (Invitrogen). Human lung (IMR-90) and skin fibroblasts (BJ) were kindly provided by Dr. Richard Gregory (Boston Children’s Hospital), and 293 cells were obtained from American Type Culture Collection. IMR-90, BJ and 293, cells were grown in DMEM (Invitrogen) supplemented with 10% FBS and 1% penicillin/streptomycin. COGN-415x cells were grown in IMEM (Invitrogen) supplemented with 1x Insulin-Transferrin-Selenium (ITS-G) (Gibco), 20% FBS and 1% penicillin/streptomycin. All cell lines were routinely tested for mycoplasma.

### Transfection

Plasmid DNA transfection was performed using Mirus Trans-IT LT1 (MIR2300) according to manufacturer’s protocol.

### Compounds

Rocaglate analog compounds, including CMLD012824 (ADR-824), were provided by Dr John Porco’s laboratory at Boston University (BU). The amidino-rocaglate (ADR) CMLD012824 was synthesized at the BU-CMD according to the reported literature procedure ^32^.

### Cell viability analysis

Cells were plated in 96-well plates at a seeding density of 4 x 10^3^ cells/well. After 24 h, cells were treated with increasing concentrations of CMLD012824 (10 nM to 10 μM). DMSO without compound served as a negative control. After 72 h incubation, cell viability was analyzed using the CellTiter-Glo Luminescent Cell Viability Assay (Promega), according to the manufacturer’s instructions. All proliferation assays were performed in biological triplicates. Drug concentrations that inhibited 50% of cell growth (IC_50_) were determined using a nonlinear regression curve fit using GraphPad Prism 6 software.

### Fluorescence-Activated Cell Sorting Analysis (FACS)

For cell cycle analysis, cells were treated with DMSO or CMLD012824 (1 nM or 5 nM). After 72 h cells were scraped and fixed in ice-cold 70% ethanol for 1 h at -20°C. After washing with ice-cold phosphate-buffered saline (PBS), cells were treated with 100 µg/mL RNase A (Sigma-Aldrich) in combination with 50 µg/ml propidium iodide (PI, BD Biosciences) for 30 min at room temperature (RT) and then kept on ice until FACS. For EdU analysis, cells were treated with DMSO or CMLD012824 (5 nM or 10 nM) for 24 h. Cells were pulsed with 10 µM of 5-ethynyl-2’-deoxyuridine (EdU) for 2 h and subsequently collected by scraping, and stained for EdU incorporation using the Click-iT EdU Alexa Fluor 647 Flow Cytometry Assay Kit (Thermo Fisher) according to manufacturer’s protocol. After EdU staining, cells were resuspended in Click-iT saponin-based permeabilization and wash reagent (Thermo Fisher) with 50 µg/mL propidium iodide (PI, BD Biosciences) and 100 µg/mL RNase A (Sigma-Aldrich) for 30 min at room temperature (RT) and then kept on ice until FACS. All samples were analyzed on an LSR Fortessa (Becton Dickinson) using FACSDiva software (Becton Dickinson). A minimum of 50,000 events was counted per sample and used for further analysis. Data were analyzed using FlowJo software.

### Apoptosis analysis

Cells were plated in 96-well plates at a seeding density of 4 x 10^3^ cells/well. After 24 h, cells were treated with increasing concentrations of CMLD012824 and analyzed using a RealTime-Glo Annexin V Apoptosis and Necrosis Assay kit (Promega JA1011) at 1 to 72 h. Annexin V binding and the loss of membrane integrity were monitored in real-time by luminescence and fluorescence respectively, according to the manufacturer’s protocol.

### Western Blotting

Cells were collected by scraping in cold PBS and lysed on ice in NP40 buffer (Invitrogen) supplemented with complete protease inhibitor cocktail (Roche), PhosSTOP phosphatase inhibitor cocktail (Roche) and PMSF (1 mM). Tumor and liver samples were prepared by washing in cold PBS, homogenizing in supplemented NP40 buffer (8k rpm, 3 sec pulses, 3-5x), and incubating on ice for 30 min. All lysates were cleared by centrifugation at 13.2k rpm for 20 min at 4°C. Protein concentrations were determined with the Biorad DC protein assay kit (Bio-Rad). Whole-cell protein lysates were resolved on 4%–12% Bis-Tris gels (Invitrogen) and transferred to nitrocellulose membranes (Bio-Rad). After blocking nonspecific binding sites for 1 h using 5% dry milk (Sigma) in Tris-buffered saline (TBS) supplemented with 0.2% Tween-20 (TBS-T), membranes were incubated overnight with primary antibody at 4°C. Chemiluminescent detection was performed with the appropriate secondary antibodies. Protein levels in western blots were quantified using ImageJ ^58^.

### Antibodies

The following antibodies were used for western blot analysis using the manufacturers’ suggested dilutions. CCNA2 (4656S), CCNE1 (4129T), CCND1 (2922S), CDK4 (12790S), CDK6 (13331S), eIF4A1 (2013S), eIF4E (9742S), eIF4G (2498S), 4E-BP1 (9644S), GAPDH (2118S), c-Jun (9165S), MYCN (51705S), c-MYC (13987S), PABP1 (4992S), cleaved PARP (5625S) [Cell Signaling].

CKS2 (37-0300), RSP19 (A304-002A-T), XRN2 (A301-103A-T) [Life Technologies]

eIF3B (A301-761A-M), eIF4A1 (ab31217), eIF4A2 (ab31218) [Abcam]

Immunoprecipitation: eIF4A1 (ab31217), PHOX2B (ab227719) [Abcam]. MYCN (51705S) [Cell Signaling]

Immunofluorescence antibodies: MYCN (51705S), PTBP1 (57246S) [Cell Signaling].

Alexa Fluor 488 goat anti-rabbit (A11008) [Abcam].

### Metabolic Labeling

Cells were incubated in L-methionine-free RPMI (A1451701) for 1 h prior to start of experiment. After methionine-free incubation, L-azidohomoalanine (Life Technologies C10102) was added according to the manufacturer’s instructions, and cells were treated with vehicle (DMSO) or CMLD012824 (1 nM, 5 nM) for 1 h. Cells were harvested by scraping in cold PBS and prepared for 1-D gel analysis using Click-IT L-azidohomoalanine protein labeling reagents (Life Technologies C10102, C10276, B10185) according to manufacturer’s instructions. Following SDS-PAGE electrophoresis and electrotransfer to nitrocellose membranes, membranes were blocked for 1 h in 5% dry milk in TBS-T (Tris-buffered saline (TBS) supplemented with 0.2% Tween-20). Biotinylated protein was visualized with NeutrAvidin Protein HRP (Thermo 31001) and chemiluminescent detection. Signal was quantified using ImageJ ^58^.

### Immunofluorescence

Cells were washed with cold PBS and fixed in 4% paraformaldehyde for 5 min, then incubated in cold 100% methanol for 5 min, and washed with cold PBS for 5 min. Cells were permeabilized with Triton X-100 0.1% for 5 minutes, washed 3x with cold PBS for x mins, and blocked for 1 h in 1% bovine serum albumin (BSA), 0.3M glycine, and 0.1% Tween-20 in PBS. Cells were incubated overnight with primary antibodies in blocking buffer, washed 3x with blocking buffer, incubated 1 h with secondary fluor-conjugated antibodies, washed 3 x with blocking buffer, and mounted on slides (25 x 75 x 1.0 mm) using Dapi Fluoromount G (OB010020). Slides were dried overnight and imaged on a Zeiss Imager Z1 Microscope.

### RT-qPCR

Total RNA was isolated with the RNAeasy Mini kit (QIAGEN) or Trizol (Thermo 15596-026) according to manufacturer’s protocol. 200 ng of purified RNA was reverse transcribed using SuperScript IV VILO master mix (Invitrogen) following the manufacturer’s protocol. Quantitative PCR was carried out using the QuantiFast SYBR Green PCR kit (Qiagen) and analyzed on an Applied Biosystems StepOne Real-Time PCR System (Life Technologies). Each individual biological sample was qPCR-amplified in technical triplicate and normalized to an internal control (input, GAPDH or other according to individual assay). Relative quantification was calculated according to the -ΔΔCt relative quantification method. Error bars indicate ± SD of three replicates. Primer sequences are available on request.

### Chromatin Immunoprecipitation (ChIP)

Dynabeads were prepared 24 h in advance by washing 50 μL beads per sample in 500 μL blocking buffer (PBS with 0.5% BSA) and incubating overnight at 4°C in 250 μL blocking buffer with 5 μg of antibody of interest or normal rabbit IgG. Bound beads were washed 3x with blocking buffer and resuspended in 100 μL blocking buffer. Cells were grown on 15 cm plates, collected by scraping in 10 mL cold PBS (1x10^8^ cells), crosslinked with 1% formaldehyde for 10 min at RT, quenched with 0.125 M glycine, washed 2x in cold PBS and flash frozen in liquid nitrogen. Cell pellets were thawed, resuspended in 5 mL LB1 (50mM HEPES-KOH pH7.5, 140mM NaCl, 1mM EDTA, 10% glycerol, 0.5% NP-40, 0.25% Triton X-100, protease inhibitor cocktail (1 tablet per 10mL)), and incubated with rotation at 4°C for 10 min. Cells were pelleted at 4k rpm for 3 min at 4°C, resuspended in LB2 (10mM Tris-HCl, pH8.0, 200mM NaCl, 1mM EDTA, 0.5mM EGTA, protease inhibitor cocktail (1 tablet per 10mL)), and incubated with rotation at 4°C for 10 min. Cells were pelleted at 4k rpm for 3 min at 4°C, resuspended in 2 mL sonication buffer (50mM HEPES pH7.5, 140mM NaCl, 1mM EDTA, 1mM EGTA, 1% Triton X-100, 0.1% Na-deoxycholate, 0.2% SDS, protease inhibitor cocktail (1 tablet per 10mL)). Cells were sonicated on ice for 30 min total time (pulse on: 30 sec, pulse off: 1 min, level 5). Sonicated samples were centrifuged at 4k rpm for 10 min at 4°C, supernatant collected and diluted with equal volume sonication buffer 2 (50mM HEPES pH7.5, 140mM NaCl, 1mM EDTA, 1mM EGTA, 1% Triton X-100, 0.1% Na-deoxycholate, protease inhibitor cocktail (1 tablet per 10mL)). 50 μL of each sample was retained for input control. 1 mL of sheared chromatin was mixed with 100 μL prepared antibody-bound beads and incubated at 4°C overnight with rotation. Beads were collected on a magnetic rack and washed 2x with sonication buffer 2 for 5 min at 4°C, 1x with sonication buffer 2 with high salt (500mM NaCl), 1x with LiCl buffer (20mM Tris pH8.0, 1mM EDTA, 250mM LiCl, 0.5% NP-40, 0.5% Na-deoxycholate), 1x with Tris-EDTA pH 8.0, and resuspended in 200 μL elution buffer (50mM Tris-HCl pH8.0, 10mM EDTA pH8.0, 1% SDS). Chromatin was eluted from beads at 65°C for 40 min with shaking, cleared on a magnetic rack, 12 μL 5M NaCl was added per sample, and samples were incubated at 65°C overnight to reverse crosslinks. The samples were then diluted 1:1 with Tris-EDTA pH 8.0, incubated with 100 μg/mL RNAse A at 37°C for 1 h, then incubated with 50 μg/mL proteinase K, 5 mM CaCl_2_ at 55°C for 30 min. DNA was extracted with 500 μL phenol:chloroform:isoamyl alcohol (EMD 516726-1SET), precipitated with 1.5 μl of GlycoBlue (Thermo AM9515), 16 μl 5M NaCl, and 1 ml 100% ice-cold ethanol at -20°C, centrifuged at 13k rpm for 20 min at 4°C, washed with 75% ethanol, and resuspended in water.

### ChIP-seq

ChIP was carried out as previously described ^5^. Purified ChIP DNA was used to prepare Illumina multiplexed sequencing libraries using the NEBNext Ultra II DNA Library Prep kit and the NEBNext Multiplex Oligos for Illumina (New England Biolabs) according to the manufacturer’s protocol. Libraries were multiplexed and sequenced using an Illumina NS500 Single-End 75bp SE75 sequencer.

### RNA Immunoprecipitation

MYCN-amplified (Kelly) and non-amplified (SK-N-AS) neuroblastoma cells were grown to 80% confluency and treated with DMSO or CMLD012824 (10 nM) for 1 h. harvested by scraping in ice-cold PBS followed by centrifugation at 500 x g for 5 min at 4°C. Cell pellets were resuspended in 1x PLB (10x PLB: 1 M KCl, 50 mM MgCl2, 100 mM HEPES-NaOH pH 7.5, 5% NP-40, Roche protease and phosphatase inhibitors (1 tab each per 10 mL)) with 200U/mL RNAsin (Promega) (3x pellet volume) and incubated on ice for 30 min. Cell lysates were centrifuged at 13k rpm for 10 min at 4°C and supernatants transferred to low-binding nuclease-free tubes. DynaBeads Protein G magnetic beads (Life Technologies 10004D) were prepared 24 h in advance by washing 2x in NT-2 buffer (5x NT-2: 250 mM Tris-HCl pH 7.4, 750 mM NaCl, 5 mM MgCl2, 0.25% NP-40) and incubating overnight with 5 μg antibody of interest or IgG control per 50 μL of beads per sample. Bound beads were washed 4x with NT-2 buffer on a magnetic rack, resuspended with 500 μL of NET-2 buffer (1x NT2 buffer supplemented with 20mM EDTA pH 8, 200U/mL Superase-In (AM2696)) plus lysate sample, and incubated overnight at 4 °C with rotation. Bound samples were washed 4x with 500 μL NT-2 buffer, resuspended in 100 μL NT-2 Buffer and divided for RNA and protein isolation. RNA samples were extracted using Trizol (Thermo 15596-026) according to manufacturer’s protocol. RT-qPCR was performed as described above. Protein samples were mixed with NuPAGE LDS Sample Buffer (Thermo NP0007) according to manufacturer’s protocol, boiled for 10 min at 95 °C, resolved on SDS-PAGE gels and analyzed by western blotting.

### Ribosome Profiling

Cells were treated with DMSO or CMLD012824 (10 nM) for 1 h. Ribosome profiling libraries were prepared from three biological replicates per cell line according to previously described methods^42^. Total RNA was extracted from matched samples using miRNeasy RNA Extraction kit (QIAGEN) and ERCC RNA Spike-In (Life Technologies 4456740) was added according to manufacturer’s instructions. RNA sequencing libraries prepared with the Illumina TruSeq stranded mRNA kit (Illumina) following the manufacturers’ instructions at the DFCI core facility. All samples were analyzed for nucleotide length and concentration (Bioanalyzer) and sequenced using an Illumina NS500 Single-End 75bp SE75 sequencer.

### PAR-CLIP

COG-N-415x PDX-derived MYCN-amplified neuroblastoma cells were grown to 80% confluency in biological triplicate on 15 cm plates, with 4-thiouridine (200 μM) (Sigma Aldrich T4509) added directly to the cell culture medium 16 h before crosslinking. Cells were treated with DMSO or CMLD012824 (10 nM) for 1 h, washed with ice cold PBS, and irradiated uncovered with 0.4 J/cm2 of 365nm UV light using Alpha Innotech AIML-26 Transilluminator. Cells were harvested by scraping and centrifugation at 2.5k rpm for 5 min at 4°C. Cell pellets were resuspended in 1x PLB (10x PLB: 1 M KCl, 50 mM MgCl2, 100 mM HEPES-NaOH pH 7.5, 5% NP-40, Roche protease and phosphatase inhibitors (1 tab each per 10 mL)) with 200U/mL RNAsin (Promega) (3x pellet volume) and incubated on ice for 30 min. Lysates were cleared by centrifugation at 12k x g for 10 min at 4°C and 10% input was saved for total mRNA sequencing library preparation. Samples were treated with RNase T1 (1 U/μl) in a water bath for 15 min at 22°C, cooled 5 min on ice, and >1 U/ul Superase-In (AM2696) was added to quench RNAse T1. DynaBeads Protein G magnetic beads (Life Technologies 10004D) were prepared 24 h in advance by washing 2x in NT-2 buffer (5x NT-2: 250 mM Tris-HCl pH 7.4, 750 mM NaCl, 5 mM MgCl2, 0.25% NP-40) and incubating overnight with 10 μg antibody (eIF4A1 ab31217) or IgG control per 100 μL of beads per sample. Samples were incubated with beads in 500 μL total volume, overnight at 4°C with rotation. Samples were washed 4x by resuspending the beads in NT-2 buffer and incubating for 5 minutes with rotation at 4°C, and resuspended in 250uL NT2 buffer, with 10 μL reserved to check IP efficiency. Samples were treated a second time with RNaseT1 (10 U/μl) at 22 °C for 20min with shaking, cooled on ice for 5 min, and washed 3x with NT-2 buffer. Bound beads were resuspended in 1 volume of dephosphorylation buffer (50 mM Tris-HCl, pH 7.9, 100 mM NaCl, 10 mM MgCl_2_, 1 mM DTT) with Calf-intestinal phosphatase (CIP) (0.5 U/μl) and incubated for 10 min at 37℃ with shaking. Beads were washed twice in 1 ml of phosphatase wash buffer (50 mM Tris-HCl, pH 7.5, 20 mM EGTA, 0.5% (v/v) NP-40), 2x in polynucleotide kinase (PNK) buffer without DTT (50 mM Tris-HCl, pH 7.5, 50 mM NaCl, 10 mM MgCl2), and resuspended in 50 μL of PNK buffer (50 mM Tris-HCl, pH 7.5, 50 mM NaCl, 10 mM MgCl2, 5 mM DTT). Samples were treated with ATP (1mM) and T4 PNK (1 U/μl) and incubated for 60 min at 37°C with shaking, washed 5x with 800 μl of PNK buffer without DTT and resuspended in 100 μl of PNK buffer without DTT. 10 μl of sample was saved for 3′ -biotin labeling for visualization using Pierce RNA 3′ End Biotinylation Kit (Life Technologies 20160) according to manufacturer’s protocol. Samples were collected on a magnetic rack, washed 3x with NT-2 buffer, resuspended in 70 μl of DEPC-treated SDS-PAGE loading buffer (NP0007) and heated for 5 min at 95°C with shaking. Beads were collected on a magnetic rack, the supernatants transferred to new tubes, resolved on a Bis-Tris 4-12% PAGE gel, and transferred to a nitrocellulose membrane (60V, 2h or 85V, 1h15min). The membrane was cut at the region determined by the 3′-biotin signal in corresponding samples using a Chemiluminescent Nucleic Acid Detection Module (Thermo 89880) according to manufacturer’s protocol. Membrane slices were treated with DNase I (5 U) in 1X DNase I buffer at 37°C for 10 min, followed by proteinase K (4 μg/ μl) digestion in PK buffer (100mM Tris-HCl pH 7.4, 50mM NaCl, 10mM EDTA) for 20 min at 37℃ with shaking, and incubated in 200 μl of PK-urea buffer (PK buffer with 7M urea) for 20 min at 37℃ with shaking. RNA was extracted with 400 μl Acid Phenol:ChCl3 (pH4.3∼4.7) and precipitated with 1.5 μl of GlycoBlue (Thermo AM9515), 40 μl NaAcO3 (pH 5.5), and 1 ml 100% ice-cold ethanol at -80°C. Samples were centrifuged at 12k x g for 60 min, washed 2x with 75% EtOH, resuspended in DEPC water, and submitted for small RNA library construction at the DFCI core facility. Total RNA was extracted from matched samples using miRNeasy RNA Extraction kit (QIAGEN). RNA sequencing libraries were processed for rRNA removal (QiaSelect) and prepared with the Illumina TruSeq stranded mRNA kit (Illumina) following the manufacturers’ instructions at DFCI core facility. All samples were analyzed for nucleotide length and concentration (Bioanalyzer) and sequenced on a NovaSeq 6000 sequencer. Two replicates per condition passed quality control (Bioanalyzer) and were used for downstream analysis.

### Sucrose gradient fractionation

MYCN-amplified (Kelly) neuroblastoma cells were grown to 80% confluency and treated with DMSO or CMLD012824 (10 nM) for 1 h. Cells were harvested by scraping in ice-cold PBS and centrifugation at 500 x g for 5 min at 4°C. Cell pellets were resuspended in 1x PLB (10x PLB: 1 M KCl, 50 mM MgCl2, 100 mM HEPES-NaOH pH 7.5, 5% NP-40, Roche protease and phosphatase inhibitors (1 tab each per 10 mL)) with 200U/mL RNAsin (Promega) (3x pellet volume) and incubated on ice for 30 min. Cell lysates were centrifuged at 13k rpm for 10 min at 4°C and supernatants transferred to low-binding nuclease-free tubes. Cellular lysates were sedimented on 10-50% sucrose gradients (containing 20 mM HEPES pH 7.5, 150 mM KOAc, 2.5 mM MgOAc, 1 mM DTT, 0.2 mM spermidine, 100 μg/mL cycloheximide) for 2 h at 40,000 g at 4 °C using an SW41 rotor (Beckman Coulter). Gradients were fractionated using Teledyne Isco Tris Peristaltic Pump and fractions were collected and pooled according to the UV trace. RNA was extracted using an equal volume of phenol:chloroform pH 6, precipitated at -20 °C overnight in 2x volume 100% EtOH, 2.7 M NaOAc, and 10 μg/mL GlycoBlue (Thermo AM9515), washed 2x in 70% EtOH and resuspended in RNase free water.

#### *In vitro* transcription

RNAs were transcribed from 1 µg of PCR-amplified templates using T7 RNA polymerase (NEB M0251S) for 2 h at 37 °C according to manufacturer’s protocol. Reactions were treated with RQ1 DNAse (Promega M6101) for 20 min at 37 °C, precipitated using 2x volume 7.5 M LiCl/50 mM EDTA at -20 °C for 1 h, washed 2x in 70% EtOH, and resuspended in RNase free water. RNAs were capped using the Vaccinia capping system (NEB M2080S) according to manufacturer’s protocol, in the presence of 20 U Superase-In (AM2696), extracted with an equal volume of phenol:chloroform pH 6, precipitated at -20 °C overnight in 2x volume 100% EtOH, 2.7 M NaOAc, and 10 μg/mL GlycoBlue Coprecipitant (Thermo AM9515), washed 2x in 70% EtOH and resuspended in RNase-free water. RNAs were capped co-transcriptionally during the T7 RNA polymerase reaction by decreasing GTP to 0.125 mM with addition of 2.5 mM cap analog (G-cap, NEB S1407S; A-cap, NEB S1406S).

#### *In vitro* translation

Rabbit reticulocyte lysates (RRL) (Promega L4960) were treated with micrococcal nuclease (NEB M0247S) and 0.8 mM CaCl_2_ for 10 min at 25 °C. Treatment was stopped with 3.2 mM EGTA. Treated RRL was incubated with 400 ng (or as indicated) T7-transcribed RNAs (5′ UTR fused to luciferase) in the presence of DMSO or CMLD012824 at the indicated concentrations, according to manufacturer’s instructions on supplemental amino acids and reaction buffer. Reactions were incubated for 1.5 hr at 30 °C and luciferase signal was measured using the Dual-Glo Luciferase Assay System (Promega E2920).

### Cloning

Endogenous 5′ UTR sequences were identified from Ensembl and RefSeq and cloned into the pcDNA4 vector backbone with Renilla or Firefly luciferase using restriction cloning. MYCN 5′ DEL and 3′ DEL deletion mutants were generated by restriction-free cloning using plasmid PCR amplification and overhang ligation using the In-Fusion Cloning Kit (Takara Bio 638910) according to manufacturer’s protocol. All 5′ UTR sequences are available in Supplementary table 2. Primer sequences are available upon request. MYCN coding sequence was inserted in frame into the pEF1a-puro vector for mammalian expression.

### Animal Studies

All procedures involving mice were guided by the DFCI Animal Care and Use Committee and performed under an IRB-approved protocol. Mouse experiments were performed using subcutaneous injections of 1x10^6^ cells into 4-6 week-old recipient female mice. NB-9464 TH-MYCN murine neuroblastoma xenografts were generated in syngeneic C57BL/6J mice, while human neuroblastoma patient-derived (COGN-415x) xenografts were generated in nude mice (NU/NU). For the first MTD study, C57BL/6J mice were treated with CMLD012824 (0.1, 0.2, 0.4 mg/kg) diluted in solvent (5.2% PEG300, 5.2% Tween-80) daily for 7 days by intraperitoneal injection. After reaching assay endpoint of 12 days, livers were excised from vehicle-treated and CMLD012824 -treated (1 mg/kg) animals for WB analysis. For the second MTD study, C57BL/6J mice bearing NB9464 tumors were randomly assigned into groups upon tumor volume reaching 100-200 mm^3^, with the volume being approximately equal between groups and treated with CMLD012824 (0.1, 0.2 mg/kg) diluted in solvent (5.2% PEG300, 5.2% Tween-80) three times per week for 30 days by intraperitoneal injection. For the efficacy study, nude mice bearing COGN-415x tumors were randomly assigned to treatment groups upon tumor volume reaching 100-200 mm^3^ and treated with 0.2 mg/kg CMLD012824 (diluted in 5.2% PEG300, 5.2% Tween-80) or vehicle control (DMSO in 5.2% PEG300, 5.2% Tween-80) three times per week for 40 days by intraperitoneal injection. Tumor size and body weight were monitored three times per week and tumor volume was calculated using the ellipsoid formula (1/2(max diameter x min diameter^2^). Once tumors reached 1000 mm^3^, the mice were euthanized according to approved animal protocols. Tumors were either fixed in 10% neutral buffered formalin, or snap frozen and stored at -80°C until further analysis. All animal experiments were conducted according to approved protocols by IACUC.

### Immunohistochemistry (IHC)

Staining was performed by Applied Pathology Systems (APS) (Shrewsbury, MA). Formalin-fixed paraffin-embedded (FFPE) tumors were stained with H&E, Cleaved Caspase 3, Ki67, or MYCN. For H&E staining, fixed tissues were dehydrated by passing through a series of ethanol solutions of increasing concentration (70-100%). Following dehydration, the tissues were cleared with xylene prior to paraffin embedding to form paraffin tissue blocks. Each FFPE block was sectioned with the thickness of 5 µm and one section was loaded to a histology glass slide. The slides were heated at 60°C for 1 h in an oven before H&E staining in the autostainer (Leica Autostainer XL). IHC was performed using a detection kit (Vector Laboratories, MP-7601) on a Dako autostainer. Paraffin sections were dewaxed, rehydrated, and subjected to antigen retrieval in Tris base buffer, pH 9.0, in a pressure cooker. Slides were blocked with BloxAll blocking buffer and 2.5% horse serum respectively prior to a 1-h incubation with anti-Ki67 antibody (Abcam, ab16667) at 1:100 dilution, anti-cleaved caspase 3 antibody (Biocare CP229A) at 1:250 dilution, or anti-MYCN antibody (Cell Signaling Technology, D4B2Y) at 1:500 dilution. Subsequently, the sections were incubated with anti-rabbit Amplifier antibody and ImmPress Excel polymer reagent sequentially before applying DAB chromogen. The slides were then counterstained with hematoxylin, followed by dehydration. Slides were scanned at APS and resulting images were analyzed with QuPath^59^.

### Computational analysis

#### Polypurine ranking analysis

Highly variable genes were identified by arraying and binning all transcripts from GSE62564 by expression level and calculating the variance coefficient using Giotto in R, as previously described ^38^. Polypurine ranking was performed in R using RefSeq sequences to extract 5′ UTR sequences and rank by [AG]_n_ motifs normalized to 5′ UTR length. 5′ UTR polypurine rank, GC rank, and additional information is available in Supplementary Table 1. Code available upon request.

#### Gene expression analysis of primary tumor dataset

RNA sequencing data in reads per million (RPM) was downloaded from Gene Expression Omnibus (GEO), accession GSE62564. Hierarchical clustering was performed in R on data pre-ranked by MYCN or c-MYC expression. Heatmap visualization of hierarchical clustering with Z-scores representing standard deviation from the mean were calculated using R package heatmap.2. The pairwise correlation matrix was generated using R package corrplot (v0.92) ^60^. Gene expression data (GSE62564) for highly variable genes from Giotto analysis was used to calculate fold changes per gene between the top and bottom 30 MYCN expressing tumors, ranked by MYCN expression. Data was divided into 10 equally sized bins based on expression (bin 1 – lowest, bin 10 – highest) or fold change (bin 1 [low] – no change, bin 10 [high] – highest positive change). Analysis of the entire data set revealed that the majority of the genes that fell within the average range of values for expression and fold-change exhibited average polypurine content and therefore were omitted from the Figure 1 (Figs. 1H-I) plots for visual clarity, with the full dataset plots included in Figure S1 (Figs. S1F, S1H). Analysis of genes correlated with eIF4A1 expression in primary tumors (GSE62564) was performed using R2 Genomics Analysis and Visualization Platform (http://r2.amc.nl).

#### ChIP-seq analysis

All ChIP-seq data were aligned using the short-read aligner Bowtie (v0.12.7)^61^ to build version GRCh37 of the human genome. To visualize ChIP-seq tracks, reads were extended by 160 bases, converted into tdf files using igvtools (v 2.2.1)^62^ and visualized in IGV ^63^. ChIP-seq peaks were detected using a peak-finding algorithm, MACS (v1.4.2)^64^ using the default P-value threshold of enrichment of 1 × 10^−5^ for all data sets. Active enhancers, ranked according to the magnitude of the H3K27ac signal, were defined as regions of ChIP-seq enrichment for H3K27ac and H3K4me1 outside of promoters. The ROSE algorithm (https://bitbucket.org/young_computation/rose)^65,66^ was used to identify super-enhancers.

Enhancers containing peaks within 12.5kb of one another were stitched together and ranked by their difference in H3K27ac signal vs input signal.

#### Ribosome profiling analysis

Raw Illumina reads from ribosome profiling and matched total RNA sequencing libraries were collapsed and adapters were trimmed using fastx_collapser from the FASTX Toolkit. Bowtie2 ^67^ was used to remove rRNA reads, TopHat2 ^68^ to align reads to the human genomes (GRCh37, GRCh38), Cufflinks v2.2.1 and Cuffdiff v2.2.1 ^69^ to extract and merge raw read counts of the biological replicates (N = 3). Samtools ^70^ was used to prepare data for genome browser visualization in IGV ^63^. Anota2seq in R was used for differential translation efficiency calculation^43^. The Anota2seq package is designed to analyze transcriptome-wide translation data (including ribosome profiling) and combines analysis of partial variance and the random variance model to normalize the input data, analyze changes in translational efficiency, and account for translational “buffering” (i.e translational efficiency changes as a function of changes in mRNA levels). The ribo-seq data was normalized against total RNA reads, which are not affected by a 1-hour ADR treatment as evident from the Anota2seq differential analysis of total RNA reads in the control-treated samples, as well as External RNA Controls Consortium (ERCC) ERCC92 synthetic RNA spike-in control sequences. A transcript with an absolute log2 fold-change ≥ 1 and a P-value ≤ 0.1 was considered significant.

#### PAR-CLIP analysis

The raw PAR-CLIP and matched total RNA sequencing reads were first processed for adapter trimming and rRNA removal as described for ribosome profiling. The data analysis was performed following the pipeline described by Jens ^46^. Identical read copies were collapsed into distinct reads and aligned to the human genome (GRCh37) with Burrows-Wheeler Alignment (BWA) v0.7.17 ^71^ allowing for up to one edit distance (mismatch, insertion or deletion). The unaligned reads were removed and aligned reads were sorted into a BAM formatted file with Samtools v1.13 ^70^. The clusters on the reference genome that eIF4A1 bound to were then identified with the pipeline-provided script that collects reads contiguously covering a section of the reference genome while screening for cross-link conversions. The identified clusters involve at least two distinct read sequences and at least one cross-link conversion. The cluster identification analysis was performed for each individual sample; also replicate consensus was taken into account, such that a cluster is reported only if it is called in both replicates. The false discovery rates (FDRs) of mapping for each sample and replicate consensus were assessed as per the pipeline by aligning the sequence reads to a decoy genome. A filter was applied to retain only those clusters between the length of 20-1000 nucleotides. This resulted in FDR ≤0.05 in every case. Clusters were annotated from Gencode (v19 annotation) and Ensembl (GRCh37.87 annotation) with genomic features (mRNA, exon, CDS, 5′ -UTR, 3′ -UTR, start codon and stop codon) and types of RNA (mRNA, lincRNA, miRNA, snoRNA, snRNA and rRNA) with intersect function in BEDTools v2.30.0 ^72^. Mapping of gene and transcript IDs of the clusters to gene symbols was carried out with biomaRt package in Bioconductor ^73^. Metagene2 in R was used to prepare metagene plots of 5′ UTR, CDS, and 3′ UTR consensus distribution of PAR-CLIP aligned reads per million (RPM) for two biological replicates per condition. For differential binding analysis, trimmed reads were aligned to human genome (GRCh37) using Bowtie v1.0.0 ^67^ and Cuffdiff v2.2.1 ^69^ was used to extract and merge raw read counts of the biological replicates (N = 2). Cuffdiff results were then used in the Anota2seq pipeline^43^ to normalize against total RNA to quantify changes in eIF4A1 binding by mRNA between vehicle and CMLD012824-treated replicates (N=2).

#### Enrichment analysis

GSEA analysis was performed on pre-ranked gene lists (polypurine analyses) and enriched pathway terms meeting a false discovery rate cutoff (FDR ≤ 0.1) were considered significant. Functional enrichment for gene sets derived from PAR-CLIP and ribosome profiling differential binding and translation regulation analyses was performed by Enrichr. All Gene ontology, Hallmarks pathway, KEGG pathway, Elsevier collection terms were ranked based on the Enrichr combined score. Enrichment of gene sets was considered significant for an adjusted P-value ≤ 0.01.

#### Motif enrichment analysis

Sequence motif enrichment analysis was performed using MEME Suite ^48^. BEDTools^72^ getfasta function was used to extract fasta sequences from 5′ UTR, CDS, and 3′ UTR regions corresponding to RNA sets of interest and Meme Motif discovery tool in MEME Suite was used to identify enriched motifs (Parameters: classic mode, site distribution = “any number of repetitions”, 0-order background model, minimum width = 4, maximum width = 16). Significant motifs were considered having E-value (P-values adjusted to motif frequency) < 0.01.

#### Integrated analysis of ribosome profiling and PAR-CLIP

COG-N-415x and Kelly cells were chosen for the comparison due to both having amplification of MYCN. The first integrative analysis identified the set of mRNAs with eIF4A1 binding clusters in ADR-824-treated PDX COG-N-415x cells (identified through PAR-CLIP) that exhibited changes in translational efficiency in ADR-824-treated Kelly cells (identified through ribo-seq). The second analysis integrated ribosome-associated mRNA changes with eIF4A1 PAR-CLIP binding changes from ADR-824-treated cells (both derived from respective Anota2Seq analyses in R).

## Statistical analysis

Statistical methods are listed in the figure legend and/or in the corresponding Methods. All quantitative analyses are expressed as the mean ± S.D. of three biological replicates, unless stated otherwise. Box plots within the violin plots defined by center lines (medians), box limits (the interquartile range between 25^th^ and 75^th^ percentiles), whiskers (minima and maxima;). Significance determined by Student’s t-test. Statistical significance for pairwise comparisons was determined using two-sided unpaired Student’s t-test, unless stated otherwise. Survival analysis was performed using the Kaplan–Meier method and differences between groups calculated by the two-sided log-rank test and the Bonferroni correction method. Tumor volume comparisons for the xenograft studies were analyzed by Welch’s test for overall efficacy analysis and Student’s t-test for individual days. Statistical comparisons of distributions of fold changes for the ribosome profiling and PAR-CLIP data were derived from Anota2Seq analysis in R, and for the ChIP-seq data from MACS. Ribosome profiling data are based on three biological replicates per condition; PAR-CLIP data based on two biological replicates per condition. ChIP–seq data are based on at least two independent experiments. GO enrichment was calculated using Fisher exact test in Enrichr.

