## Supplementary Figures for "The MYCN 5′ UTR as a therapeutic target in neuroblastoma"

### Supplemental Figure 1

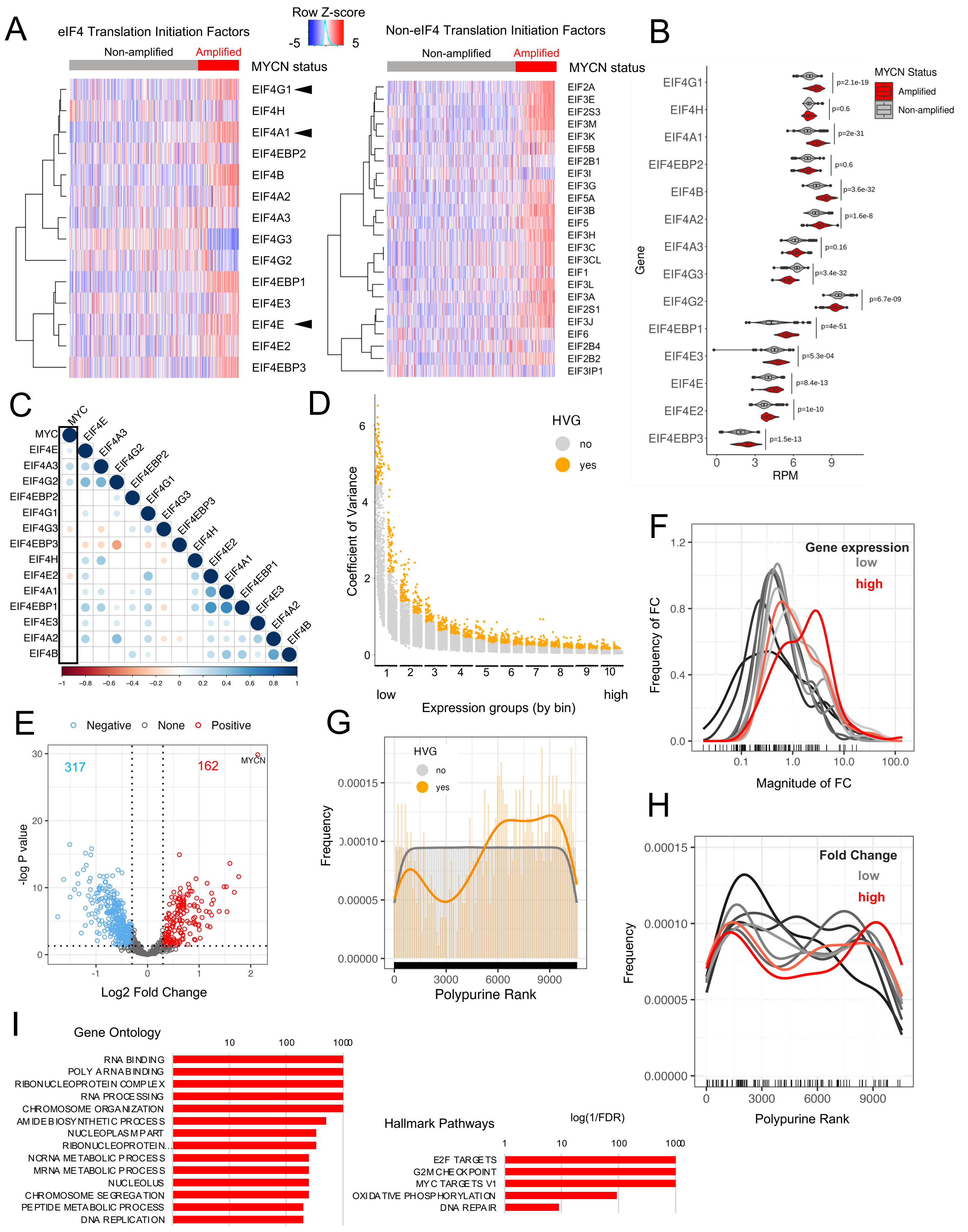

**Figure S1. Translation initiation machinery genes are correlated with MYCN but not c-MYC overexpression. A**. Hierarchical clustering of translation initiation factor gene expression in primary NB tumors (n=498, derived from GSE62564) grouped by annotated MYCN amplification status. Z-score mean ± S.D. Bar above heatmap represents corresponding MYCN expression level in log reads per million (log2 RPM). **B.** Violin plots showing expression of the indicated initiation factors in tumors with amplified and nonamplified MYCN, as depicted in A. Box plots within the violin plots defined by center lines (medians), box limits (the interquartile range between 25th and 75th percentiles), whiskers (minima and maxima). Significance determined by Student's t-test. **C.** Correlogram of c-MYC and translation initiation factor gene expression in MYCN-nonamplified NBs (n = 401, GSE62564). Circles represent Spearman's rank correlation coefficients, P <0.01. Color code represents positive correlations in blue, negative correlations in orange-red. **D.** Dot plot showing highly variable genes (HVG) identified from gene expression data in primary NB tumors (n=498, GSE62564). Variance was determined by arraying and binning all genes by expression level and calculating the variance coefficient for each group, which was then converted into a z-score. Significant HVGs with a z-score > 0.1 are depicted in gold. **E.** Volcano plot showing changes in expression of highly variable genes (HVGs) in tumors with lowest and highest MYCN expression levels (n=30 each, GSE62564). Y-axis shows significance of variance in log P-value, with horizontal line representing cutoff of $P < 0.01$. X-axis shows log2 fold change. Upregulated genes (>2-fold change) are shown in red, downregulated genes in blue, Student's t-test, $P < 0.01$. **F.** Fold change distributions of highly variable genes (HVGs) in tumors from lowest (black, bins 1 and 2) to highest (red, bins 9 and 10) MYCN expression levels (n=30 each, GSE62564) (Student's t-test, P < 0.01). X-axis shows the magnitude and Y-axis shows the frequency of fold change (FC) in expression. **G**. Contour plot showing the polypurine rank distribution of highly variable genes (HVGs). **H.** Polypurine rank distribution of the highly variable upregulated genes from lowest (black, bins 1 and 2) to highest (red, bins 7 and 8) fold change. (Student's t-test, P < 0.01). **I**. Functional enrichment analysis of the top 25% polypurine-rich genes ranked by MYCN expression in NB primary tumors (n=498). FDR <0.1.

### Supplemental Figure 2

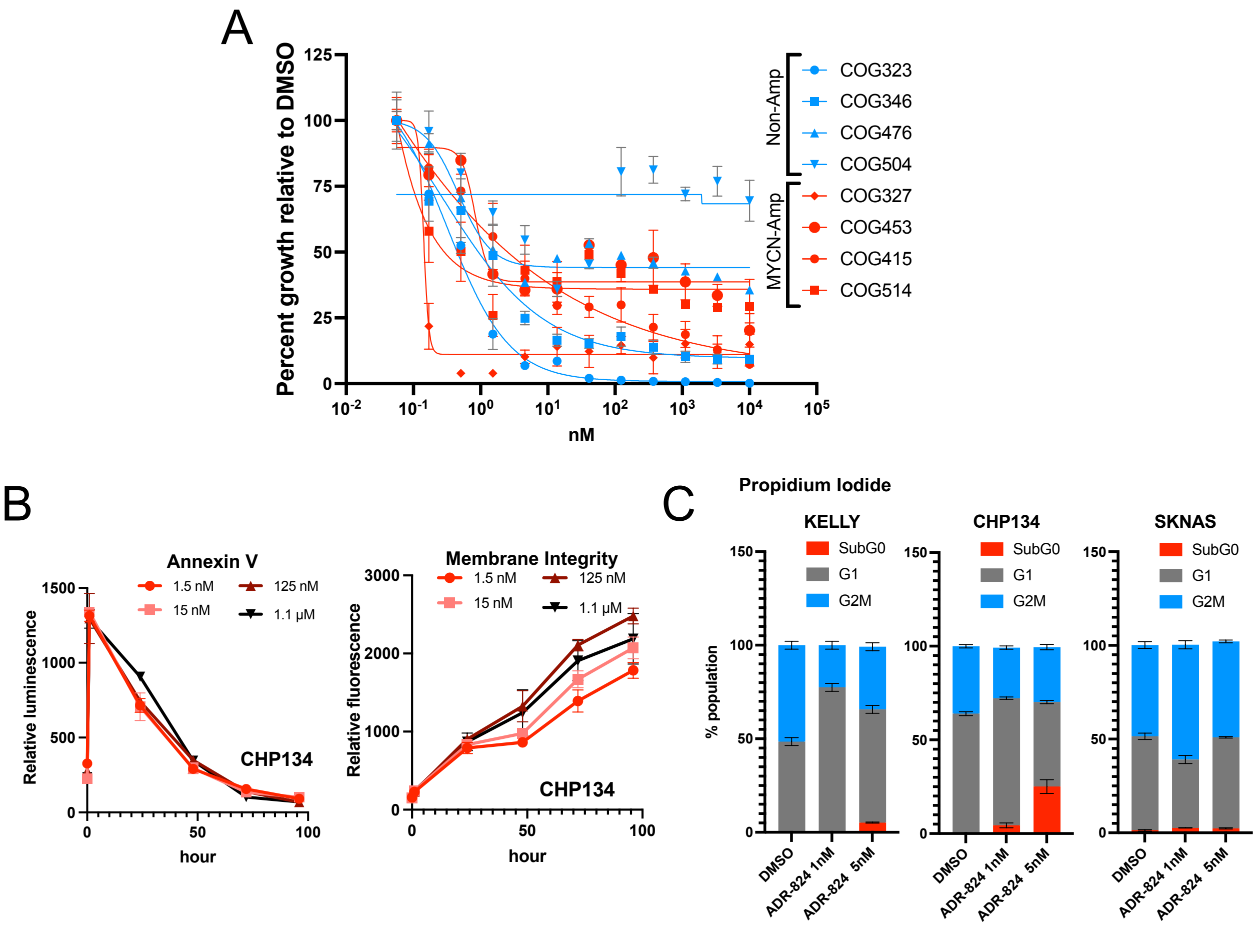


**Figure S2. CMLD012824 leads to differential cytotoxicity in NB**.

**A.** Cell viability of MYCN amplified (red) and non-amplified (blue) human PDX-derived NB cells, treated with varying concentrations of CMLD012824 (ADR-824) for 72 h. Percent cell viability relative to DMSO is shown. Data represent mean ± S.D., n = 3 replicates. **B.** Annexin V (left) and membrane integrity (right) analysis at the indicated times following treatment with the indicated doses of ADR-824 in MYCN-amplified (CHP-134) cells. Data represent mean ± S.D., n = 3 replicates. **B.** Quantification of propidium iodide incorporation in the indicated NB cells 72 h post treatment with ADR-824 at the indicated doses. Data represent ± S.D., between n = 3 biological replicates.

### Supplemental Figure 3

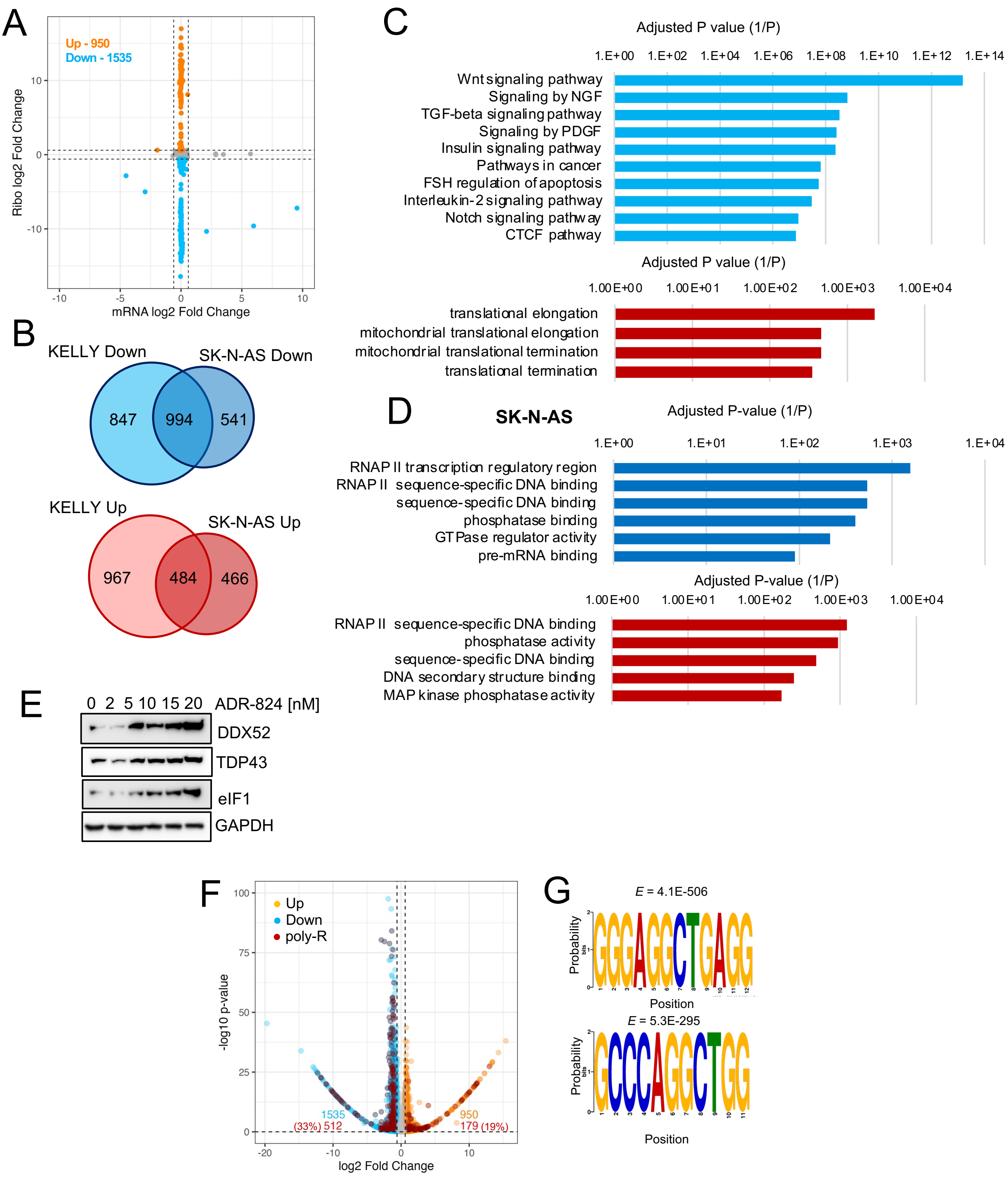

**Figure S3. ADR-824-mediated inhibition targets unique subsets of long, poly-purine-rich mRNAs in MYCN-amplified versus nonamplified NB cells.**

**A.** Scatter plot of total vs. ribosome associated mRNA changes in ADR-824-treated MYCN-nonamplified (SK-N-AS) NB cells (10 nM, 1 h), with axes showing log2 fold change as determined by Anota2seq (P <0.1) (See Methods). Dotted black lines indicate cut-offs of 1.5-fold change in total mRNA (x-axis) and ribosome occupancy (y-axis) of ADR-824-treated biological replicates (n=3) compared to vehicle control (n=3). Data points are color coded according to ribosome occupancy, with corresponding total numbers shown on the plot (Upregulated shown in gold, downregulated shown in blue). **B.** Venn diagram representing the overlap in translationally regulated MYCN-amplified (Kelly) and nonamplified (SK-N-AS) gene sets**.** Down: >1.5-fold decrease, Up: >1.5-fold increase (*P* <0.1). **C.** Functional enrichment of common downregulated (top, blue) and upregulated (bottom, red) processes in MYCN-amplified and non-amplified NB cells identified by ribosome profiling analysis [Down: >1.5-fold decrease, Up: >1.5-fold increase (*P* <0.1)]. **D.** Functional enrichment of unique downregulated processes in non-amplified (SK-N-AS) cells [Down: >1.5-fold decrease (*P* <0.1)]. Axis depicts log of inverse P-value in **C, D**. **E.** WB analysis of representative unique upregulated mRNAs in MYCN-amplified (Kelly) cells, with GAPDH as control. **F.** Volcano plot of translationally regulated mRNAs in ADR-824-treated MYCN-nonamplified (SK-N-AS) NB cells, showing translational efficiency changes as determined by Anota2seq (>1.5-fold change, *P* <0.1). Polypurine-rich (top 25%) mRNAs are highlighted in red, with percentages indicated on the plot. **G.** Motif enrichment analysis showing top motifs in the downregulated subset, trained against a background list of unregulated transcripts. **H**. Ranking of super-enhancer associated genes in MYCN-amplified (Kelly) NB cells based on histone 3 lysine 27 acetylation (H3K27ac) signal, plotted in order of increasing super-enhancer rank (MYCN=1, highest). Translational efficiency changes and polypurine ranking were calculated as in (4C). Upregulated mRNAs are shown in gold, downregulated in blue, >1.5-fold change Polypurine-rich (top 25%) mRNAs are highlighted in red. Right: quantification of polypurine-rich (top 25%) and polypurine-poor (bottom 25%) in the downregulated mRNA subset.

### Supplemental Figure 4

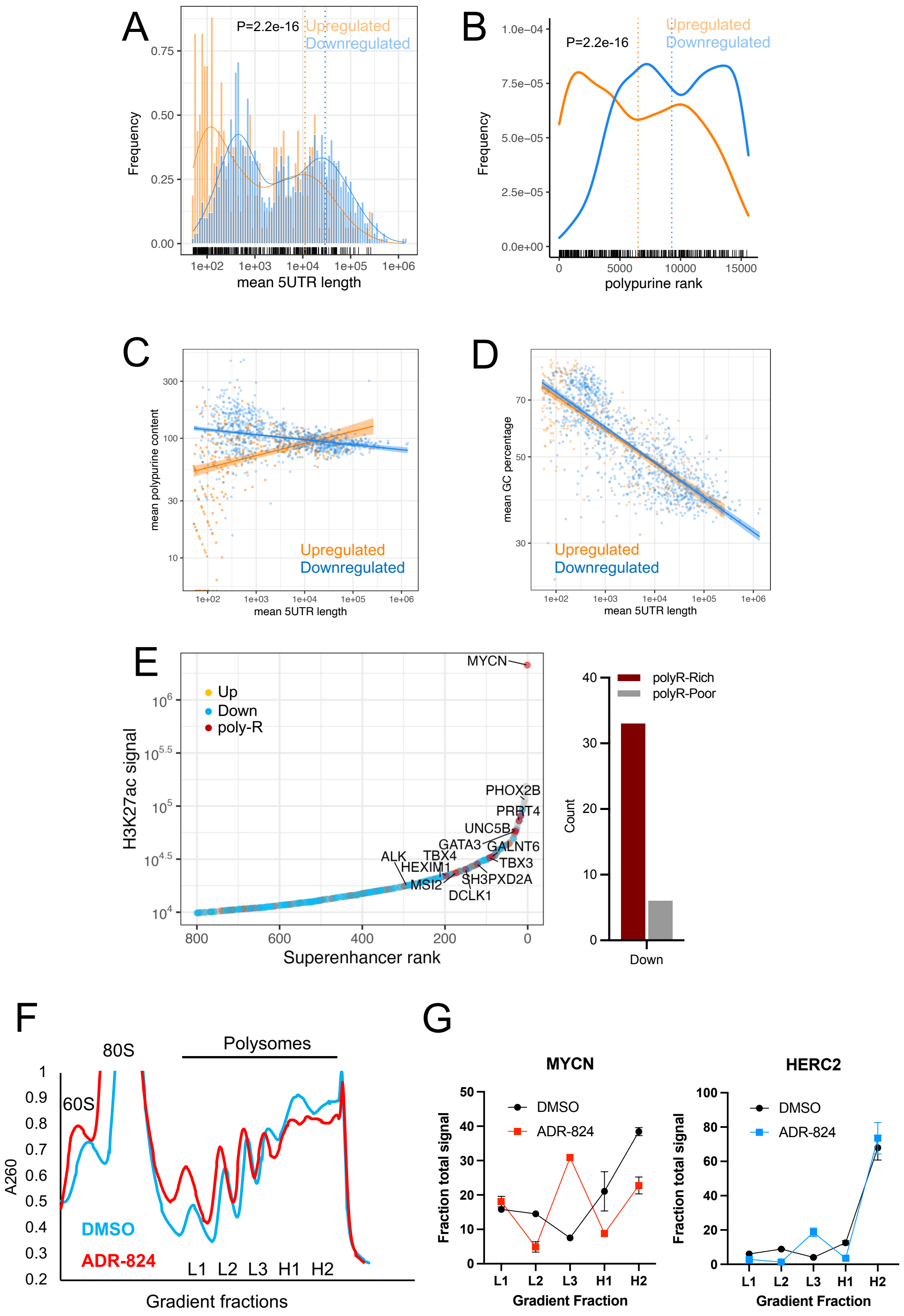

**Figure S4. ADR-824-mediated inhibition alters ribosome occupancy of poly-purine-rich mRNAs in MYCN-amplified NB cells. A.** Contour plot showing 5′ UTR length distribution of translationally regulated transcripts in (gold: upregulated, blue: downregulated) in ADR-824-treated MYCN-nonamplified (SK-N-AS) NB cells. *P* <2.2e-16 (up- vs. down-regulated mRNAs, >2-fold change, Student's t-test). **B**. Contour plot of polypurine rank distribution of translationally regulated transcripts in nonamplified (SK-N-AS) NB cells. *P* <2.2e-16; Student's t-test. **C, D.** Scatter plot of polypurine content (**C**) and GC content (**D**) by 5′ UTR length in upregulated (gold) versus downregulated (blue) mRNAs in nonamplified cells. Loess regression analysis is shown in corresponding colors, shaded regions represent 95% confidence intervals. **E.** $A_{260}$ absorbance signal of RNA in sucrose gradient sedimentation fractions. Kelly cells were treated for 1 h with vehicle or ADR-824 (10 nM), lysed and fractionated on 10-50% sucrose gradients by ultracentrifugation. The positions of 80S, light (L1-3) and heavy (H1-2) polysomes are indicated on the plot. Results show one representative experiment. **F.** RT-qPCR analysis of MYCN and HERC2 polypurine-poor control mRNA distribution in polysome fractions after vehicle or ADR-824 treatment in MYCN-amplified (Kelly) NB cells. Light polysomes: L1-3; heavy polysomes: H1-2. Signal was calculated by $2^{-\Delta\Delta Ct}$ method, normalized to total RNA in gradient. Data represent mean ± S.D., n = 3 replicates.

### Supplemental Figure 5

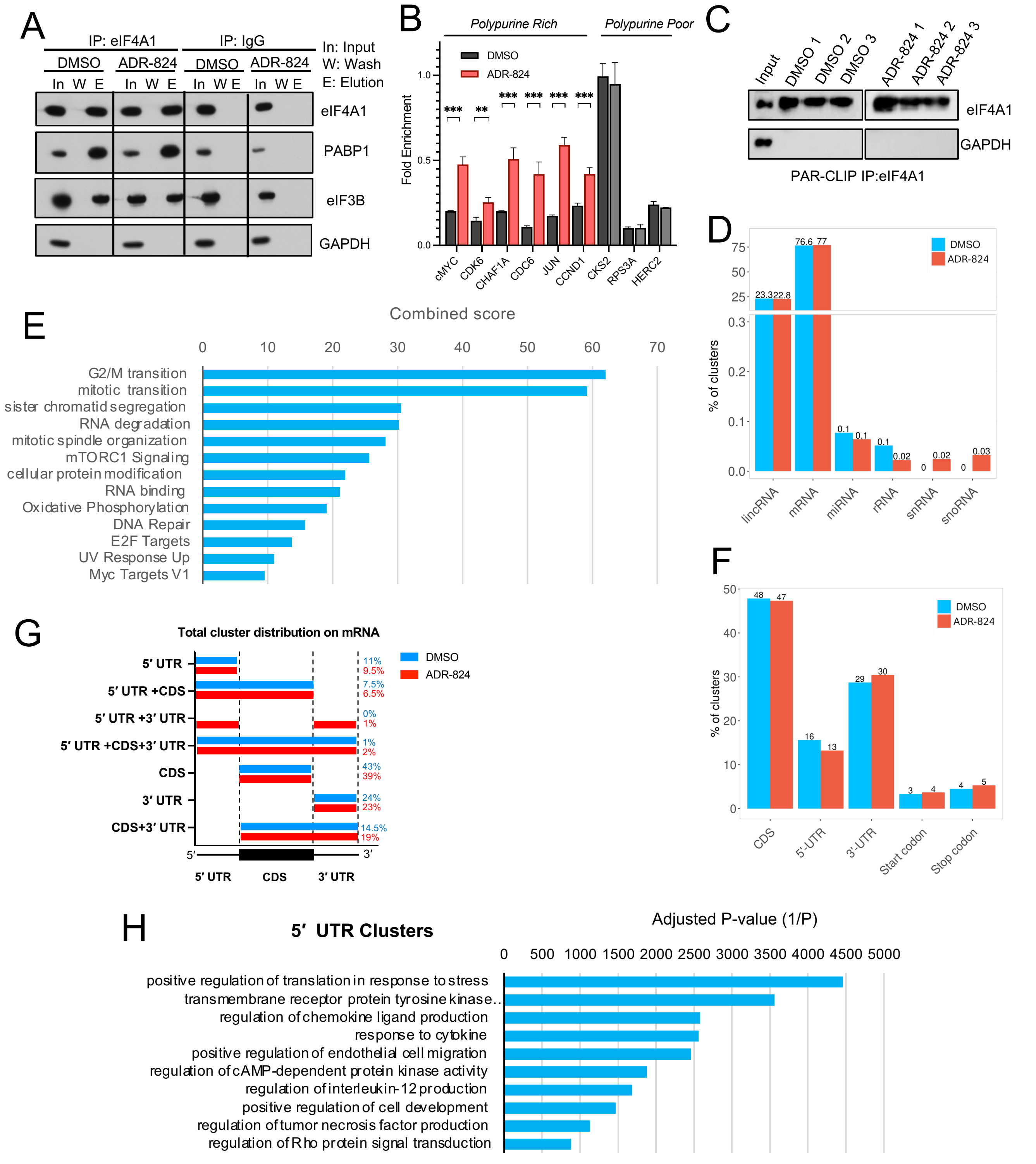

**Figure S5. eIF4A1 binds along the full length of mRNAs and other classes of RNAs. A.** WB analysis of eIF4A1 and the indicated proteins (with GAPDH used as a control) in immunoprecipitates of eIF4A1 or IgG from DMSO- and ADR-824-treated MYCN-amplified (Kelly) cell lysates. **B.** RT-qPCR analysis of 5′ UTR polypurine-rich and -poor mRNAs bound to eIF4A1 immunoprecipitated from MYCN-nonamplified (SK-N-AS) cell lysates following treatment with DMSO or ADR-824 (10 nM x 1 h). Data represent mean ± S.D., n = 3 replicates. **p value < 0.001, ***p value < 0.0001, Student's t-test. **C**. WB analysis of immunoprecipitation (IP) of eIF4A1 after PAR-CLIP, with GAPDH serving as loading and crosslinking (negative) control. **D.** Distribution of eIF4A1-binding clusters in DMSO- and ADR-824-treated cells that map to the indicated classes of RNAs. **E.** Functional enrichment analysis of eIF4A1-bound clusters from ADR-824-treated cells. Combined score indicates P-value derived from Fisher's exact test in Enrichr ($P < 0.001$ cutoff). **F**. Distribution of eIF4A1-binding clusters that map to the indicated mRNA regions. **G**. Distribution of eIF4A1-binding clusters to the indicated regions of mRNAs in ADR-824-treated cells**.** Average percentages out of the total unique mRNAs per replicate (DMSO, n = 826, 1833; ADR, n= 13128, 12593) indicated on right. **H**. Functional enrichment analysis of eIF4A1-binding clusters from ADR-824-treated cells that map to the 5′ UTRs. Combined score indicates adjusted p value derived from Fisher's exact test in Enrichr ($P < 0.001$ cutoff).

### Supplemental Figure 6

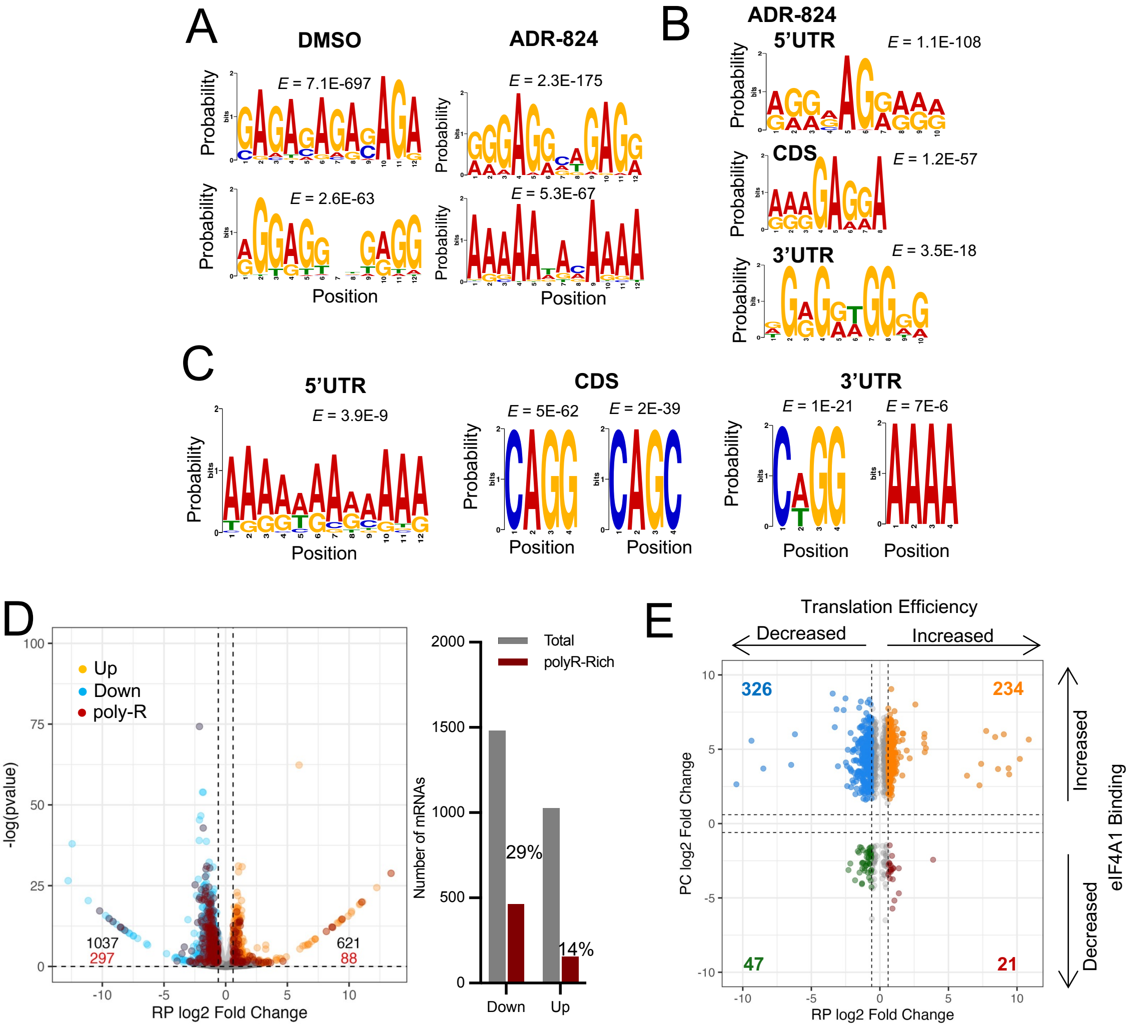

**Figure S6. eIF4A1 binds to polypurine motifs along the full length of mRNAs.**

**A.** Top motifs identified in eIF4A1-binding clusters in DMSO- and ADR-824-treated cells. E-values adjusted to motif frequency. **B.** Additional top discovered motifs from ADR-824-treated eIF4A1-binding clusters that map to the indicated mRNA regions. **C.** Top motifs identified in eIF4A1 clusters that map to the indicated mRNA regions in ADR-824-treated cells, trained against the DMSO-treated background. E-values adjusted to motif frequency are shown. **D.** Left, Volcano plot of translationally regulated mRNAs (identified through ribo-seq) that correspond to eIF4A1-bound RNAs (identified through PAR-CLIP) in ADR-824-treated cells. The x-axis shows the ribosome profiling (RP) translational efficiency and y-axis, statistical significance, as determined by Anota2seq (>1.5-fold change, $P$ <0.1). Polypurine-rich (top 25%) mRNAs are highlighted in red. Right: Quantification of polypurine-rich mRNAs with >2-fold change in translational efficiency, in down- and upregulated ribo-seq subsets. **E.** Scatter plot of ribosome-associated mRNA changes (x-axis) versus eIF4A1 PAR-CLIP binding changes (y-axis) from ADR-824-treated cells, with axes showing log2 fold change as determined by Anota2seq ($P$ <0.1). Data points are colored according to changes in ribosome occupancy and eIF4A1 binding (>1.5-fold change in both) with corresponding numbers shown on the plot. Dotted black lines indicate cut-offs for ADR-824-treated compared to DMSO-treated biological replicates (1.5-fold change, n=2 each).

### Supplemental Figure 7

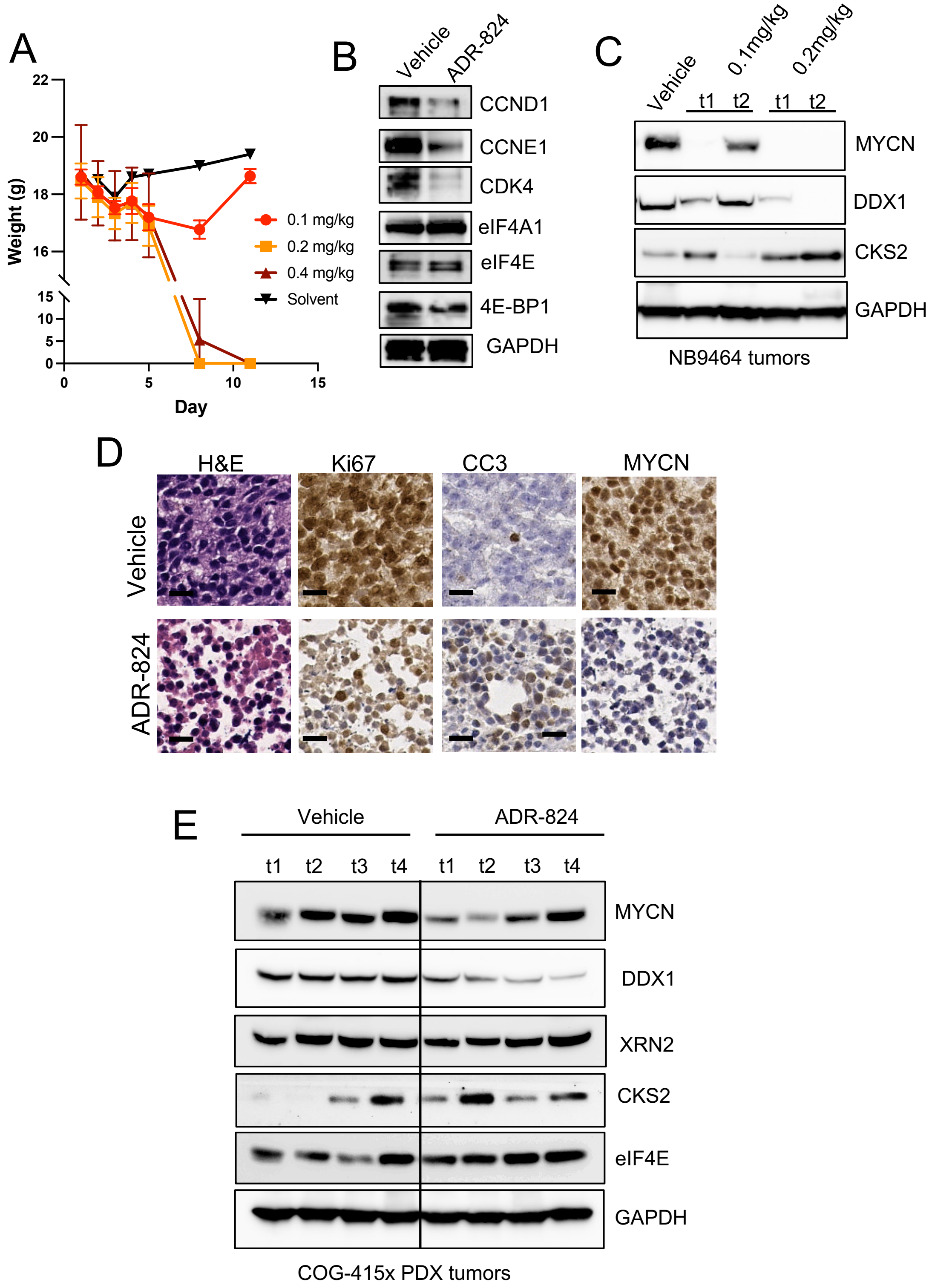

**Figure S7. Establishment of maximum tolerated dose of ADR-824 in murine models.**

**A.** Serial weights of C57BL/6J non-tumor-bearing mice treated daily for 5 days with the indicated doses of ADR-824 or vehicle. Weights are shown in grams ± S.D. n = 2 per ADR-824 treated, n=1 for vehicle-treated. Weight of zero indicates animal death. **B.** WB analysis of proliferation and cell cycle proteins (CCND1, CCNE1, CDK4) and translation initiation regulatory proteins (eIF4A1, eIF4E, 4E-BP1) in liver tissue of vehicle and ADR-824 (0.1mg/kg) mice in (A). GAPDH serves as a loading control. **C.** WB analysis of polypurine-rich (MYCN, DDX1), -poor (CKS2), and GAPDH control proteins in NB-9464 PDX tumors in Fig 7A. t# – tumor designation in group. **D.** H&E and IHC analyses of additional representative vehicle (top) and ADR-824-treated (bottom) COGN-415x-derived tumors. Scale bar represents 100 µm. **E.** WB analysis of polypurine-rich (MYCN, DDX1), -poor (CSK2, XRN2), and control (eIF4E, GAPDH) proteins in COG-N-415x PDX tumors in Fig 7A. t# – tumor designation in group.
